# Disulfide bonds are required for cell division, cell envelope biogenesis and antibiotic resistance proteins in mycobacteria

**DOI:** 10.1101/2025.01.27.635063

**Authors:** Adrian Mejia-Santana, Rebecca Collins, Emma H. Doud, Cristina Landeta

## Abstract

Mycobacteria, including *Mycobacterium tuberculosis*—the etiological agent of tuberculosis—have a unique cell envelope critical for their survival and resistance. The cell envelope’s assembly and maintenance influence permeability, making it a key target against multidrug-resistant strains. Disulfide bond (DSB) formation is crucial for the folding of cell envelope proteins. The DSB pathway in mycobacteria includes two enzymes, DsbA and VKOR, required for survival. Using bioinformatics and cysteine profiling proteomics, we identified cell envelope proteins dependent on DSBs. We validated via *in vivo* alkylation that key proteins like LamA (MmpS3), PstP, LpqW, and EmbB rely on DSBs for stability. Furthermore, chemical inhibition of VKOR results in phenotypes similar to those of Δ*vkor*. Thus, targeting DsbA-VKOR systems could compromise both cell division and mycomembrane integrity. These findings emphasize the potential of DSB inhibition as a novel strategy to combat mycobacterial infections.

## Introduction

*Mycobacterium tuberculosis*, the causative agent of tuberculosis, causes 1.5 million deaths worldwide every year, and the emergence of multidrug-resistant strains poses a serious threat to global health^1–3^. Additionally, nontuberculous mycobacteria (NTM) are an underrecognized source of lung disease and the incidence of NTM disease has been surging worldwide becoming an emerging public health problem^4^. Mycobacteria have an intricate cell wall that represents a formidable barrier to drug entry and is a major contributor to resistance. Mycobacterial cell envelope is unusual in that constitutes mycobacterial peptidoglycan covalently linked to arabinogalactan, which is also covalently bound to mycolic acids^5^. This complex structure is also known as mycolyl-arabinogalactan-peptidoglycan complex^5^. The bacterial cell envelope is the interface with the host and is critical to reacting to immune factors, tolerating antibiotic treatment, and adapting to the variable host environment^6^. Thus, targeting the assembly and maintenance of the mycobacterial cell envelope represents a strategy against multidrug-resistant bacteria.

Oxidative protein folding is a promising novel target because disulfide bonds (DSBs) play a critical role in folding multiple exported proteins in the bacterial cell envelope. We and others have proposed to target DSB-forming enzymes in pathogens as a new strategy to combat the antibiotic resistance crisis^7–10^. Likewise, inhibition of DSB formation re-sensitizes multidrug-resistant clinical isolates of gram-negative bacteria including *Escherichia coli, Klebsiella pneumoniae,* and *Stenotrophomonas maltophilia* to currently available antibiotics^11,12^.

In *E. coli* DSB formation involves two enzymes. First, a periplasmic oxidoreductase, DsbA, enables appropriate folding of proteins by removing a pair of electrons^13^. Then, the membrane protein DsbB regenerates DsbA’s activity by transferring electrons to quinones^14–16^, either ubiquinone aerobically or menaquinone (aka Vitamin K2) anaerobically^17^. Most proteobacteria have DsbAB homologs, conversely actinobacteria such as Mycobacteria, use an enzyme called <u>V</u>itamin <u>K</u> ep<u>o</u>xide reductase (VKOR) instead of DsbB (Fig. 1a)^18,19^. Bacterial VKOR is a homolog of the human VKORc1 enzyme^19,20^, yet bacterial and mammalian VKOR proteins display divergent mechanisms of electron transfer to quinones^21,22^.

**Figure 1.**
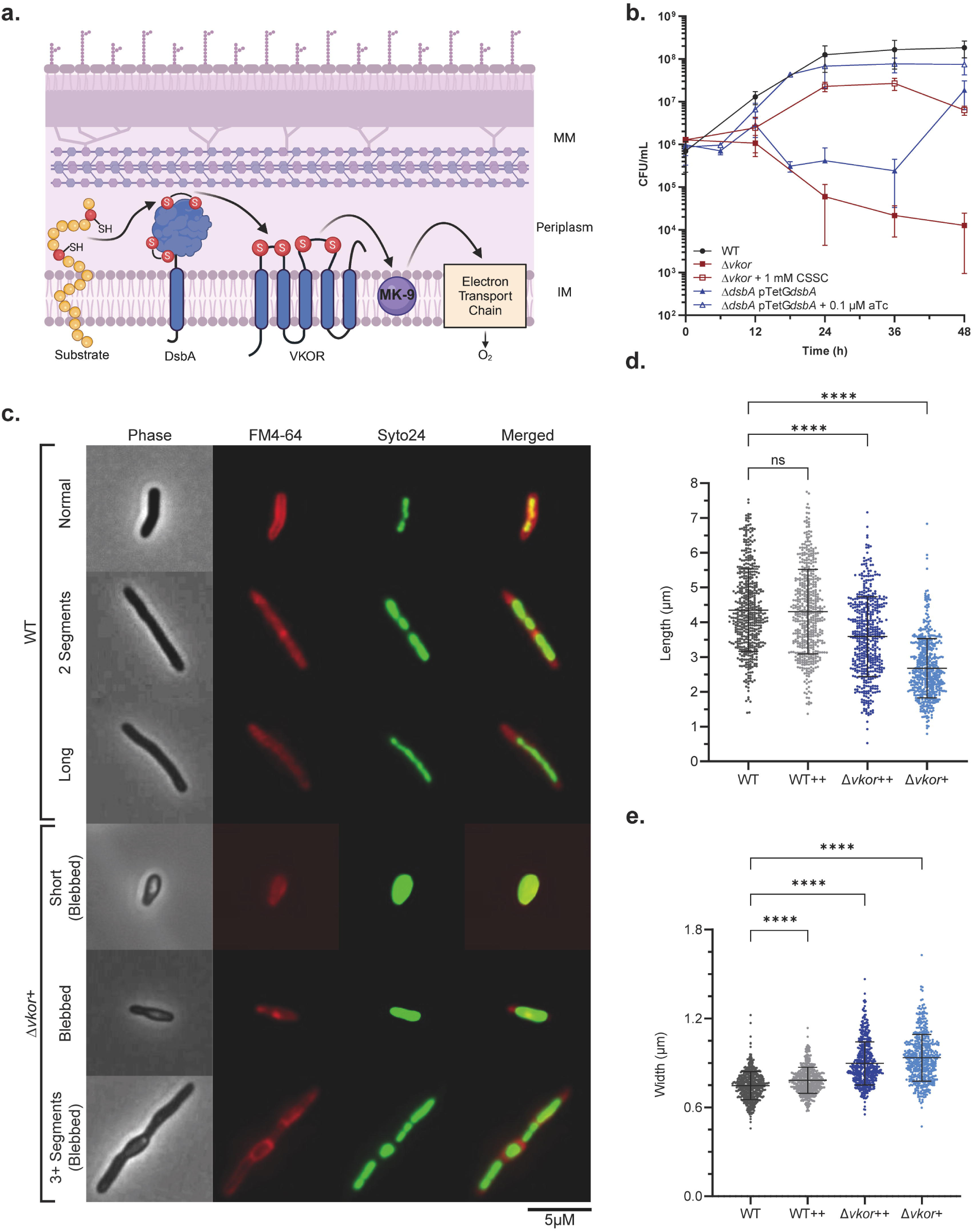
Lack of DSB formation causes growth and cell division defects in *M. smegmatis*. **a,** DsbA oxidizes secreted proteins and becomes reduced. VKOR regenerates reduced DsbA by transferring the electrons to menaquinone. Black arrows represent the flow of electrons, IM: inner membrane, MM: mycomembrane, MK-9: menaquinone-9. **b,** *M. smegmatis* Δ*dsbA* and Δ*vkor* survival is affected in modified 7H9 broth. Δ*dsbA* was supplemented with 100 nM of aTc to induce Ms*dsbA* expression and Δ*vkor* was supplemented with 1 mM cystine (CSSC). Data represents the average ±SD of at least two independent experiments. **c,** WT and Δ*vkor* cells supplemented with 1 mM (++) or 100 µM (+) cystine, were fluorescently stained with 50 nM Syto24 to stain nucleic acids and 0.6 µg/mL FM4-64 to stain the membrane. Representative images of the phenotypes are shown. **d-e,** Cell dimensions were measured from samples obtained from at least three independent experiments using FIJI (https://fiji.sc/). Data represents the average ±SD. Cell counts included: WT (n=495), WT++ (n=505), Δ*vkor*++ (n=498), Δ*vkor*+ (n=500). Statistical tests were done using Kruskal-Wallis multiple comparisons. p-values are depicted in GP style: ≤0.0001 (****), 0.0002 (***), 0.021 (**), 0.0332 (*), and non-significant (ns).

While a detailed comparison of the mechanism of electron transfer between *M. tuberculosis* VKOR and *E. coli* DsbA revealed strong parallels to the *E. coli* DsbB and DsbA pair^23^, the mycobacterial proteins have unique features. First, *M. tuberculosis* DsbA contains an additional pair of cysteines that form a DSB and is anchored to the plasma membrane by a single transmembrane (TM) segment^24–26^. Second, VKOR contains five TM helices as opposed to four found in *E. coli* DsbB^27^ or human VKORc1^28^. Finally, the *M. tuberculosis* predicted exported proteome identifies thousands of proteins that contain even numbers of cysteine residues, suggesting that DSBs might be important for the folding and function of extracytoplasmic proteins^19^. Nevertheless, substrates of the DsbA-VKOR pathway remain largely unknown.

In this work, we investigated the essential proteins that require oxidative protein folding in mycobacteria by using bioinformatics, cysteine-profiling proteomics, and genetic and biochemical approaches. We identified 787 proteins that were decreased in *M. smegmatis* Δ*vkor* grown under semi-permissive conditions, while 398 proteins increased. Furthermore, cysteine enrichment proteomics revealed 745 oxidized cysteines corresponding to 381 secreted proteins, which are potential substrates of DsbA. Using *in vivo* alkylation, we validated that *M. tuberculosis* LamA (MmpS3), PstP, LpqW, and EmbB require one to two DSBs, when expressed in *M. smegmatis,* a model mycobacterial species^29^. We also demonstrated that the stability of LamA, PstP, LpqW, and EmbB is compromised in the absence of DSBs in Δ*vkor* and PstP cysteine mutants exhibit impaired growth. Finally, we show as a proof of concept that chemical inhibition of VKOR phenocopies the Δ*vkor* mutant.

## Results

### Disulfide bonds are important for cell division and septation in *M. smegmatis*

*M. tuberculosis* DsbA and VKOR essentiality was discovered through transposon mutagenesis^30^. Similarly, *M. smegmatis* lacking the *vkor* gene could be generated when a small molecule oxidant, cystine (oxidized cysteine), is supplemented in the media^31^ (Fig.1b). When the Δ*vkor* is grown without cystine there is a rapid decline in the population after 12 h (Fig. 1b), indicating the need for an oxidant to regenerate DsbA’s activity. Contrastingly, efforts to delete the *dsbA* homolog in *M. smegmatis* were unsuccessful with the addition of cystine^31^. Most likely cystine cannot oxidize secreted proteins directly but is more efficient in re-oxidizing DsbA in the Δ*vkor*^31^. The deletion of *dsbA* is only possible in a merodiploid strain expressing *dsbA* from a regulatable promoter^31^. This conditionally lethal mutant is unable to grow in the absence of the DsbA-inducer, anhydrotetracycline (aTc) (Fig.1b). When this strain is grown without inducer, however, suppressors arise after 36 h (Fig.1b). Presumably these suppressors are mutations that disrupt TetR regulation given their high frequency^32^.

We sought to determine the underlying mechanism DsbA-VKOR have on mycobacterial survival by first identifying morphological defects in *M. smegmatis* Δ*vkor* (Fig. 1c). When Δ*vkor* is grown under limiting concentration of cystine (0.1 mM, Δ*vkor* +), the cells are smaller and wider compared to wildtype (WT, Fig. 1d,e). These cells displayed membrane blebs indicative of cell wall defects and eventually, they lyse (Fig. 1c). While Δ*vkor* grows well under higher concentrations of cystine (1 mM, Δ*vkor* ++) there is still a population of smaller cells, indicating that cystine is not as effective at oxidizing DsbA as VKOR is (Fig. 1d). Additionally, we observe that 63% of cells of the Δ*vkor* under low cystine display blebbed morphology and 82% are undivided (Supplementary Fig. 1a-c, Δ*vkor* +) compared to the 1% of blebbed and 27% of undivided cells observed in WT. The Δ*vkor* grown under semi-permissive conditions also displays 27% of cells with three or more segments compared to the 1% observed in WT (Supplementary Fig. 1b). Altogether these data indicate that a decrease in DSB formation causes a defect in cell division and septation.

### *In-silico* analysis and cysteine profiling proteomics reveal potential substrates of mycobacterial DsbA

Based on the morphologies seen in Δ*vkor*, we hypothesized that protein(s) involved in cell division and/or cell envelope biogenesis requiring DSBs are the reason for the DsbA-VKOR essentiality. Thus, we performed an *in-silico* analysis using the previously identified 625 essential genes in *M. tuberculosis*^30^. Since DSB formation occurs in exported proteins, we selected the essential proteins with a predicted signal sequence or TM segments. We narrowed down the list based on proteins experimentally confirmed to be secreted^33^ and harboring extracellular cysteines. Out of 625 proteins, 19 met these requirements and represent potential DsbA substrates (Fig 2a). These proteins are conserved across all mycobacteria as well as their cysteine residues (Supplementary Table 1 and Supplementary Fig. 2). The 19 candidates are mainly involved in cell envelope biogenesis and maintenance.

**Figure 2.**
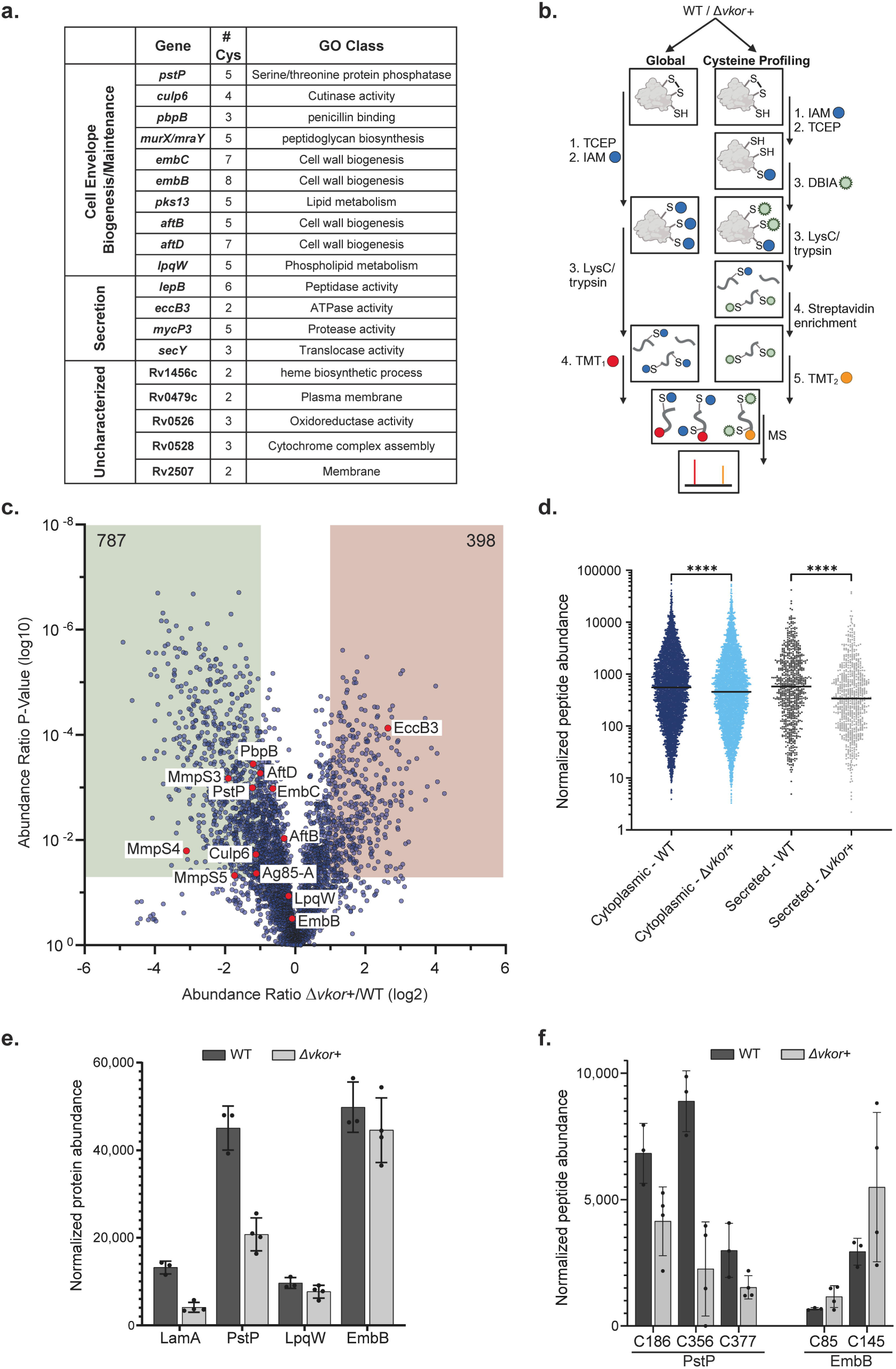
*In silico* and proteomic analyses reveal candidate DsbA substrates. **a,** Bioinformatic analysis to mine essential secreted genes identifies 19 candidate DsbA substrates (see Supplementary Table 1). **b,** Global and cysteine profiling proteomics approach to identify potential DsbA substrates (see text for further details). TCEP, tris(2-carboxyethyl)phosphine; IAM, iodoacetamide; DBIA, desthiobiotin-iodoacetamide; TMT, tandem mass tag; MS, mass spectrometry. **c,** Volcano plot of protein abundance ratios of Δ*vkor*+ compared to WT (see Supplementary Table 2). Dowregulated (green) or upregulated (pink) proteins with log2 ratios of ±1 and p-value≤0.05 are highlighted. Red dots indicate proteins analyzed or discussed in this work (see Supplementary Table 4). **d,** Biotinylated-cysteine containing peptides were sorted by the presence of TM segments or signal sequences and their normalized abundance was plotted, data represents the median abundance. Statistical test was done using Kruskal-Wallis multiple comparisons of the mean. p-values are depicted in GP style: ≤0.0001 (****), 0.0002 (***), 0.021 (**), 0.0332 (*), and non-significant (ns). **e,** Normalized abundance of four proteins of interest detected by global proteomics are shown. Data represents the average ±SD of at least three independent replicas. **f,** Abundance of cysteine containing peptides of two proteins of interest detected by cysteine profiling proteomics. Data represents the average ±SD of at least three independent replicas.

Parallel to the *M. tuberculosis in silico* analysis, we performed global and cysteine derivatization and enrichment proteomics using *M. smegmatis* WT and Δ*vkor* (grown with 0.5 mM cystine) to identify oxidized cysteines. For this, we adapted a method previously developed to globally quantify modified cysteines in mouse tissues^34,35^. This approach relies on blocking reduced cysteines with an alkylating molecule, iodoacetamide (IAM), which binds irreversibly to free thiols (Fig. 2b). Oxidized (including disulfide-bonded) cysteines are then reduced with tris(2-carboxyethyl)phosphine (TCEP) and newly reduced cysteines are labeled with desthiobiotin-iodoacetamide (DBIA) (Fig. 2b). Proteins are then digested with trypsin and desthiobiotinylated peptides are enriched using streptavidin beads for mass spectrometry analysis, this enrichment step is crucial for achieving broader coverage of the cysteine proteome^34^. In parallel, the sample is also run for proteome wide analysis to determine changes in protein abundance (Fig. 2b).

Through the global approach, we identified 4,021 unique proteins (Fig. 2c and Supplementary Table 2), of which 787 were significantly decreased in the Δ*vkor+* proteome (abundance ratio≤0.5 and p-value≤0.05) while 398 proteins were significantly increased (abundance ratio≥2 and p-value≤0.05). We then mined the protein dataset for the presence of TM segments or signal sequences and found that 38% (298) of the downregulated and 17% (66) of the upregulated proteins are predicted to be secreted (Supplementary Table 2). We mapped protein-protein interactions within these two protein groups to determine their relationships, as some proteins may change in response to the absence of DSBs in other proteins. We found 22 clusters in the downregulated and 12 in the upregulated (Supplementary Fig. 3), one key cluster includes cell division among the downregulated proteins. This cluster has three essential proteins including PstP, FtsW and PknB (Supplementary Table 3).

Additionally, we identified 6,328 desthiobiotinylated (DB) cysteine modified peptides through the enrichment approach (Supplementary Table 2). These peptides correspond to 5,607 unique cysteine residues and 2,545 unique proteins that experienced cysteine oxidation. From these 87% (4862 peptides) are cytoplasmic while 13% (745 peptides) belong to proteins with predicted TM or signal sequences (Supplementary Table 2). The median normalized abundance of DB-peptides in Δ*vkor* is 41% and 17% lower than WT in secreted and cytoplasmic proteins, respectively (Fig. 2d). Although the reduction of some of these peptides is due to a decrease in the overall protein abundance (Supplementary Table 2). Lastly, the 745 DB-peptides correspond to 381 secreted proteins that represent potential substrates of DsbA. Among these, 25 are essential, with some also found to be downregulated, such as PstP, AftD and LppZ while others, including EccE3 and EccB3 are upregulated (Supplementary Table 3).

To validate potential DsbA-VKOR substrates found in our *in silico* and proteomics screens, we focused on four proteins shown to be crucial for cell division and antibiotic resistance in *M. tuberculosis*, two of these appeared in our proteomics approach and two in our bioinformatic analysis. First, we selected MmpS3 or LamA (for loss of <u>a</u>symmetry <u>m</u>utant A), which along with other MmpS homologs (MmpS4 and Mmps5), was significantly less abundant (Fig 2de); and the bioinformatic analysis also identified a MmpS homolog, an essential protein of unknown function, Rv2507 (Fig 2a). Mycobacterial MmpS proteins are usually not essential for survival^36^ and they all harbor a conserved pair of cysteines (Supplementary Fig. 3). LamA has a critical role in inhibiting cell wall synthesis at the new poles^37^—creating the characteristic asymmetric growth of mycobacteria that leads to heterogeneity in the cell population, and consequently impacts antibiotic resistance. Second, PstP is the only serine/threonine phosphatase in *M. tuberculosis* involved in growth rate, cell length, cell division, and cell wall metabolism^38–41^. PstP appeared decreased in Δ*vkor* (Fig 2e), its first three cysteines were enriched in the cysteine profiling indicating oxidation and these peptides were less enriched in Δ*vkor+* (Fig. 2f and Supplementary Table 3). Third, LpqW, a lipoprotein that regulates phosphatidylinositol mannosides, lipomannan, and lipoarabinomannan required for mycomembrane biogenesis and thus an intrinsic contributor of antibiotic resistance^42,43^. LpqW is a candidate of the *in silico* analysis but shows no decrease in the proteome wide analysis (Fig 2e) and none of its cysteines were enriched. Fourth, EmbB, an arabinosyltransferase that catalyzes branching of arabinogalactan required for cell envelope biogenesis^44^ and one of the targets of ethambutol, a first-line antituberculosis drug^45^. EmbB appeared in the *in silico* search but show no decrease in the global proteomics (Fig 2e). The cysteines of EmbB were found in the oxidized cysteine enrichment in both WT and Δ*vkor*, and in fact Δ*vkor* displayed higher enrichment perhaps due to more accessibility to trypsin digestion in the mutant (Fig 2f).

### *M. tuberculosis* LamA, PstP, LpqW, and EmbB require DsbA-VKOR for protein stability

To determine whether these selected proteins require folding through DsbA-VKOR we fused them to a 3X-FLAG tag at the carboxy terminus and ectopically expressed them in both *M. smegmatis* WT and Δ*vkor* mutant supplemented with permissive (Δ*vkor++*) and semi-permissive (Δ*vkor+*) concentrations of cystine. We found that *Mt* LamA is indeed 91% less abundant in the *vkor* mutant grown under semi-permissive conditions (0.5 mM cystine) relative to WT (Fig. 3a). Even under permissive growth (1 mM cystine) the same decrease in abundance, ∼88%, is observed. Similarly, *Mt* PstP is ∼44% lower in Δ*vkor* grown with 1 mM cystine and ∼95% lower grown with 0.4 mM cystine (Fig. 3a). *Mt* LpqW displays a ∼46% and ∼80%reduction in the Δ*vkor* grown with high and low cystine, respectively (Fig. 3a). As for *Mt* EmbB, we observe ∼62% and ∼67% reduction in the *vkor* grown under 1 mM and 0.5 mM cystine, respectively (Fig. 3a). Importantly protein degradation was not observed in the Δ*vkor* mutant, when expressing a cytoplasmic protein involved in peptidoglycan synthesis, *Mt* MurF (Fig. 3a). Thus, these results suggest that *Mt* LamA, *Mt* PstP, *Mt* LpqW, and *Mt* EmbB may be misfolded and degraded due to the limited amount of DSBs in Δ*vkor*.

**Figure 3.**
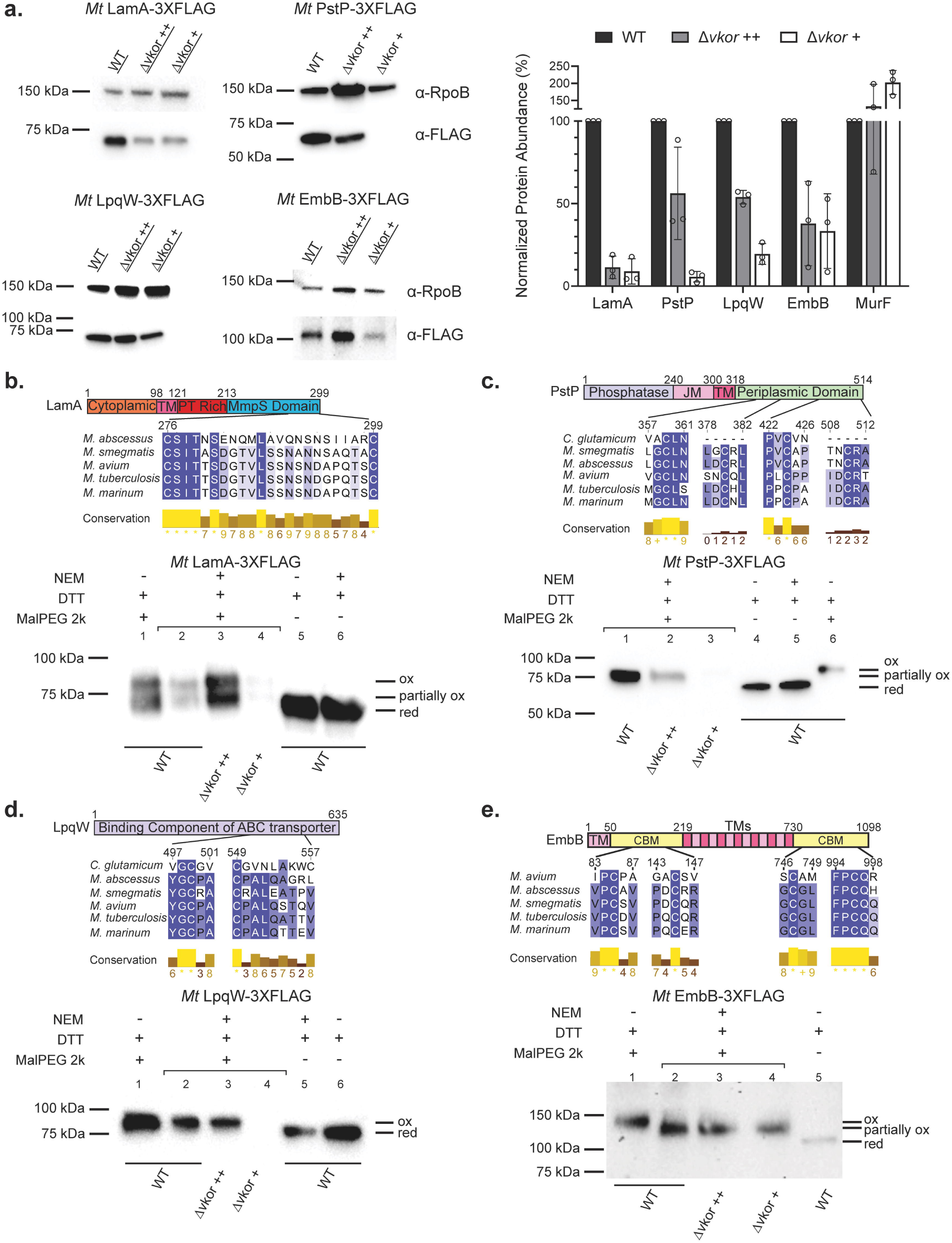
*M. tuberculosis* LamA, PstP, LpqW and EmbB are substrates of DsbA/VKOR. **a,** *Mt* LamA, *Mt* PstP, *Mt* LpqW, and *Mt* EmbB require DSBs for stability. *M. tuberculosis* proteins were fused to a 3X-FLAG tag at their carboxy termini and expressed in *M. smegmatis* WT and Δ*vkor* supplemented with 1 mM (++) or 0.4-0.5 mM cystine (+). Cells were grown at 37°C in the presence of 5 nM (EmbB) or 200 nM aTc for 36 h (PstP and LpqW), or 200 nM aTc for 18 h (LamA). Proteins were precipitated from cell lysates, quantified, and reduced before being separated by SDS-PAGE. Protein abundance was determined using α-RpoB as a loading control. Representative images are shown. Data represent average±SD of three independent experiments. **b,** *Mt* LamA harbors one DSB. Top, multiple sequence alignment was done using clustal omega^83^ and Jalview to visualize it (https://www.jalview.org/). Bottom, *M. smegmatis* WT and Δ*vkor* expressing *Mt* LamA were grown and induced as indicated in a. Experimental protein samples (indicated with a bracket) were differentially alkylated by treating them with 20 mM NEM to block free thiols. Disulfide-bonded cysteines were then reduced with 100 mM DTT, and new thiols were alkylated with 12.5 mM MalPEG2k. Controls were treated with 100 mM DTT and then alkylated with either 20 mM NEM or 12.5 mM MalPEG2k. Δ*vkor* samples were loaded in excess to be able to observe alkylated bands. Western blotting using α-FLAG antibody was used to detect LamA. Immunoblot is a representative image of at least three independent experiments. **c,** *Mt* PstP harbors two DSBs. Similar as in b. **d,** *Mt* LpqW harbors one DSB. Similar as in b. **e,** *Mt* EmbB harbors two DSBs. Similar as in b.

### *M. tuberculosis* LamA, PstP, LpqW, and EmbB harbor disulfide bonds

To investigate whether these proteins harbor DSBs when the DsbA-VKOR system is intact, we performed *in vivo* differential alkylation to derivatize the reduced cysteines. This method consists of precipitating proteins from cell lysates and blocking irreversibly all free thiols with a light alkylating agent, NEM (N-ethylmaleimide, 0.125 kDa). Disulfide-bonded cysteines are then reduced with dithiothreitol (DTT), and all nascent free thiols are labeled with a second heavy alkylating agent, MalPEG2k (α-[3-(3-Maleimido-1-oxopropyl)amino]propyl-ω-methoxy, polyoxyethylene, 2 kDa). Thus, the molecular weight of the proteins with DSBs would show an increase in size of 2 kDa per cysteine, while non-disulfide bonded cysteines would have a marginal size change that can be identified via SDS-PAGE and western blotting. The advantage of this approach is that disulfide-bonded cysteines in a given protein display an increase in molecular weight as opposed to no shift in conventional alkylation labeling with only one heavy alkylating molecule.

*Mt* LamA is predicted to have a single TM helix and a periplasmic MmpS domain harboring the two conserved cysteines (Supplementary Fig. 5a). *In vivo* differential alkylation revealed that *Mt* LamA, of a predicted size of 33.8 kDa, runs in a dimeric state of approx. ∼68 kDa (Fig. 3b, lanes 5 and 6) and accumulates in an oxidized state (+4 kDa) in WT and Δ*vkor*, albeit at lower abundance in the mutant (∼72 kDa. Fig. 3b, lane 2-4). A partially oxidized state with one cysteine labeled is also observed for WT and the Δ*vkor* mutant (Fig. 3b, lanes 2-4). This state could be an intermediate with a cysteine inaccessible to MalPEG2k due to steric hindrance since it is also observed in the alkylated control rather than a DsbA-LamA intermediate. We used a heavier alkylating agent (MalPEG5k) to corroborate this finding and observed a similar result (Supplementary Fig. 6a). Here, we also observed that Δ*vkor* with high and low cystine concentration have a small amount of the reduced form (Supplementary Fig. 6a, lane 1 vs. 2-3), consistent with the reduced form of *Mt* LamA being unstable and degraded. Collectively, these results suggest the presence of one DSB in *Mt* LamA.

Similarly, we determined the presence of DSBs in *Mt* PstP. Its topology prediction indicates that one cysteine is located in the cytoplasm near one of the catalytic residues (Asp188) and four cysteines are located in the periplasm (Supplementary Fig. 5b). *In vivo* differential alkylation indicates that *Mt* PstP is accumulated in a partially oxidized state in WT (Fig. 3c, lanes 1; Supplementary Fig. 6b, lane 2). This state indicates that one cysteine is in reduced state hence alkylated with the light agent (+0.125 kDa) while the other four cysteines were labeled with the heavy agent (+8 kDa) after being reduced. Thus, *Mt* PstP with a predicted molecular weight of ∼56.6 kDa (Fig. 3c, lane 4; Supplementary Fig. 6b lane 1) migrates slightly faster (∼64.6 kDa, Fig. 3c, lane 1; Supplementary Fig. 6b lane 2) than the control with five labeled cysteines (∼66.6 kDa. Fig. 3c, lane 6; Supplementary Fig. 6b lane 3). The partially oxidized state is also seen in the Δ*vkor* although at lower abundance dependent on cystine concentration (Fig. 3c, lane 2-3). No reduced form (∼57.22 kDa) can be observed in the Δ*vkor*. We performed *in vivo* alkylation to corroborate this finding labelling only with MalPEG2k. We observe that *Mt* PstP displays a small shift of ∼2 kDa (Supplementary Fig. 6c, lane 1) above the non-alkylated control (Supplementary Fig. 6c, lane 4), while the fully labeled control (Supplementary Fig. 6c, lane 7) has a ∼10 kDa shift as expected thus, indicating only one cysteine of PstP is reduced. Altogether, these results suggest the presence of two DSBs in *Mt* PstP.

Subsequently, *Mt* LpqW contains five cysteines, but two of these are located within the signal sequence and one belongs to the conserved lipobox motif^46^ (Supplementary Figure 5c). Therefore, only two cysteines are potentially forming a DSB. Both *in vivo* differential alkylation with MalPEG2k and MalPEG5k show that LpqW (∼66 kDa) is present in an oxidized state in WT and Δ*vkor* under high cystine concentration (Fig. 3d, lanes 2 and 3; Supplementary Fig. 6d, lanes 2 and 3) and less of this form is observed in Δ*vkor* with low cystine thus suggesting the presence of one DSB.

Lastly, *Mt* EmbB (∼121 kDa) harbors fifteen TM segments and eight cysteines but only four of these are periplasmic (Supplementary Fig. 5d). *In vivo* differential alkylation with MalPEG2k revealed a partial shift indicating a partially oxidized state (Fig. 3e, lane 1 vs 2). The half shift suggests that half of its cysteines are disulfide-bonded indicating that *Mt* EmbB contains two DSBs. This partially oxidized form is also observed in Δ*vkor* albeit in less abundance with low cystine. Similarly to the other substrates, no reduced form can be detected (Fig. 3e, lane 2 vs 3-4).

In sum, we present evidence that *Mt* LamA, *Mt* PstP, *Mt* LpqW and *Mt* EmbB harbor one, two, one, and two DSBs, respectively in *M. smegmatis* and the folded form of these proteins decreases in the *vkor* mutant in a dose-dependent manner to cystine concentration.

### *M. tuberculosis* PstP requires two consecutive and essential disulfide bonds

Proteins containing two DSBs, such as *Mt* PstP and *Mt* EmbB, have two rearrangement options; either cysteines are disulfide bonded consecutively as they appear after being translocated to the periplasm^47^ or they are rearranged by an isomerase between nonconsecutive cysteines after being oxidized by DsbA^48^. We selected *Mt* PstP to determine the consecutiveness of the two DSBs given the size, multiple TMs and cysteines in *Mt* EmbB. To do this, we generated cysteine to serine changes in *Mt* PstP and compared their protein abundance when alkylated. We observed that when the last two cysteines are mutated there is ∼90% decrease in protein compared to WT (Fig. 4a, lane 4 and 5), while serine substitutions in the first two periplasmic cysteines were below the limit of detection (Fig. 4a, lane 8 and 9). Noteworthily, the cytoplasmic cysteine, Cys189, mutant also displays ∼80% decrease perhaps due to its proximity to the catalytic site (Fig. 4a). Our data suggests that *Mt* PstP harbors two consecutive DSBs and the first DSB between Cys359 and Cys380 provides slightly more stability than the DSB between Cys424 and Cys510.

**Figure 4.**
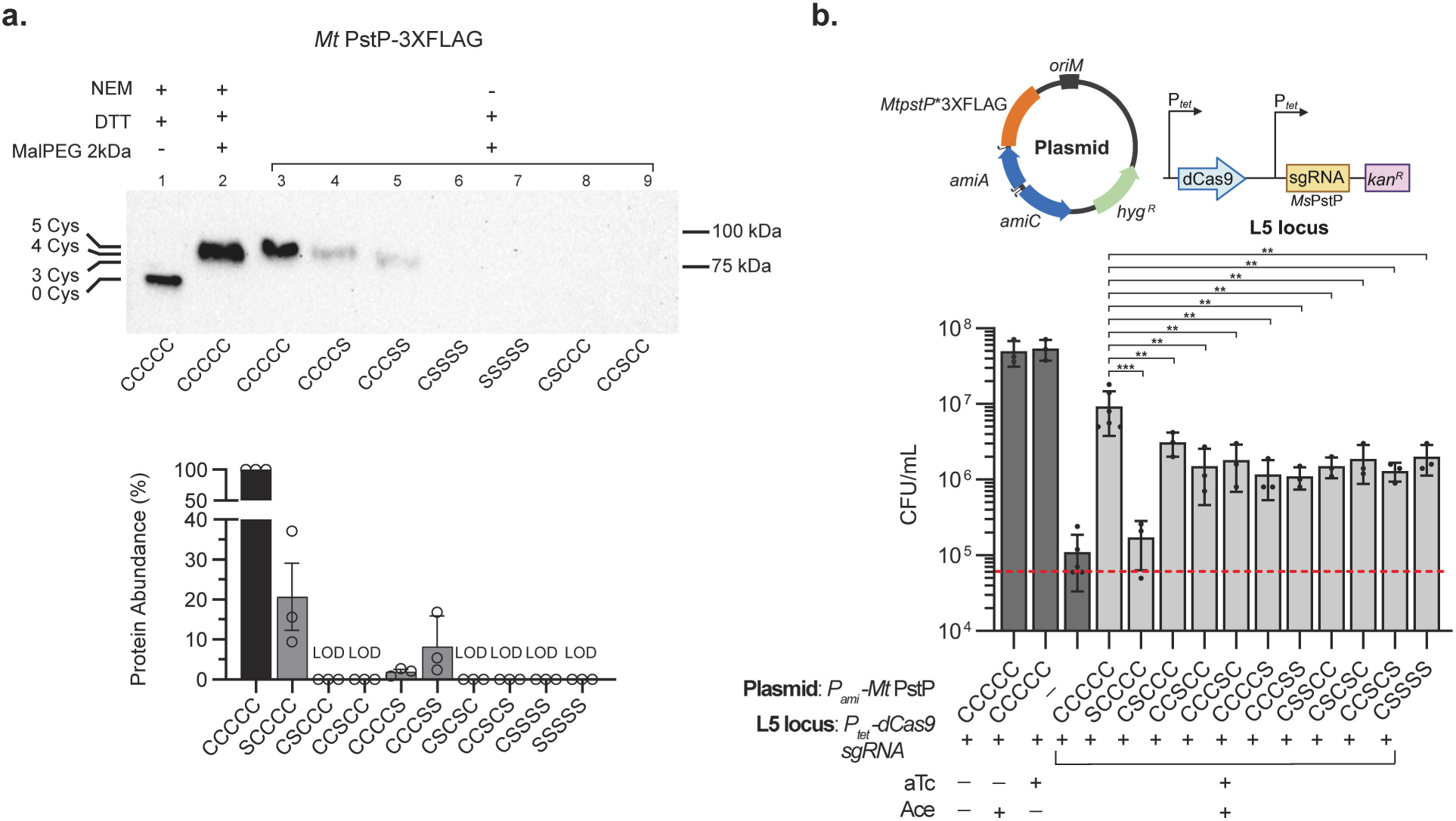
*M. tuberculosis* PstP harbors two essential consecutive DSBs. **a,** The first DSB between Cys359 and Cys380 of PstP provides more stability than the DSB between Cys424 and Cys510. Cells were grown at 37°C for 36 h in the presence of 200 nM aTc. Proteins were precipitated, reduced with 100 mM DTT and alkylated with 12.5 mM MalPEG2k. Controls were either reduced or diferentially alkylated as indicated in Fig. 2. Immunoblot is a representative image of three independent experiments. Protein abundance was determined using reduced samples and α-RpoB was used as a loading control. Data represent average±SD of three independent experiments. LOD: below the limit of detection. Cysteines are indicated in order: C189, C359, C380, C424, and C510. **b,** DSBs in PstP are required for survival. *M. smegmatis pstP* was silenced using CRISPRi while an ectopic copy of *M. tuberculosis pstP*, either WT or cysteine mutants, was used to rescue the knockdown growth. Cells were inoculated to an OD_600_ of 0.01 (CFU/mL indicated as red dotted line) in 7H9 broth supplemented with 400 nM aTc and 25 µM of acetamide (Ace), and incubated at 37°C for 24 h to enumerate bacteria. Statistical tests were done using one-way ANOVA multiple comparisons. p-values are depicted in GP style: ≤0.0001 (****), 0.0002 (***), 0.021 (**), 0.0332 (*), and non-significant (ns).

To test the role of each of the two DSBs in *Mt* PstP we generated a knockdown using a previously developed CRISPRi system for mycobacteria^47^. We inserted *tet*-regulated dCas9 and sgRNA targeting Ms*pstP* at L5 site and complemented it with an ectopic copy of *Mt pstP* under the control of the acetamidase promoter (P*_ami_*). Induction of CRISPRi with 400 nM aTc silences chromosomal *Ms pstP*, which prevents viability (Fig. 4b, aTc) and causes a characteristic blebbing phenotype in cells^39^ (Supplementary Fig. 7a). Growth and morphology can then be restored by expressing *MtpstP* with 25 µM acetamide (Fig. 4b, Supplementary Fig. 7a). We replaced the *MtpstP* copy with cysteine mutant alleles to determine the essentiality of the two DSBs. Even though all periplasmic cysteine mutants were unstable as shown above (Supplementary Fig. 7b), they affected to some extent the growth of the *pstP* knockdown (Fig. 4b). This reduction in growth is comparable to the reduction observed when the non-essential periplasmic domain is deleted^38^. The cytoplasmic cysteine displayed more impact on survival, possibly because it is near the catalytic domain. Furthermore, both mutants in either DSB were equally affected in growth despite their differences in protein degradation (Fig. 4b). However, the characteristic blebbing phenotype observed in Δ*pstP* mutant can be observed as slightly more pronounced when lacking the first (21%) or both (28%) rather than the second (16%) DSB (Supplementary Fig. 7a). Altogether, the two consecutive DSBs in *Mt* PstP are partially required for survival.

### Inhibition of mycobacterial VKOR phenocopies **Δ***vkor* mutant

We determined as a proof of concept whether inhibition of VKOR in *M. smegmatis* can have the same effect as the Δ*vkor* on essential proteins. We used bromindione (BR), an oral anticoagulant, which is a modest but, so far, one of our best inhibitors of mycobacterial VKOR^22,48^. When *M. smegmatis* is grown in 750 µM BR, the Minimal inhibitory concentration (MIC), displays slower growth compared to the WT control (Fig. 5a) and this condition phenocopies the Δ*vkor* grown on high cystine (Fig. 1b). The cell length and morphology (Fig. 5b. Supplementary Fig. 8a-b) of BR-treated cells also resemble the Δ*vkor* grown on high cystine concentration. Furthermore, at 36 h of growth BR-treated cells have 90% reduction of disulfide bonded *Mt* PstP compared to the vehicle control (Fig. 5c), and the *Mt* PstP stability is dose dependent (Fig. 5c). The effect of BR in *Mt* PstP levels is comparable to the Δ*vkor* grown on low cystine concentration.

**Figure 5.**
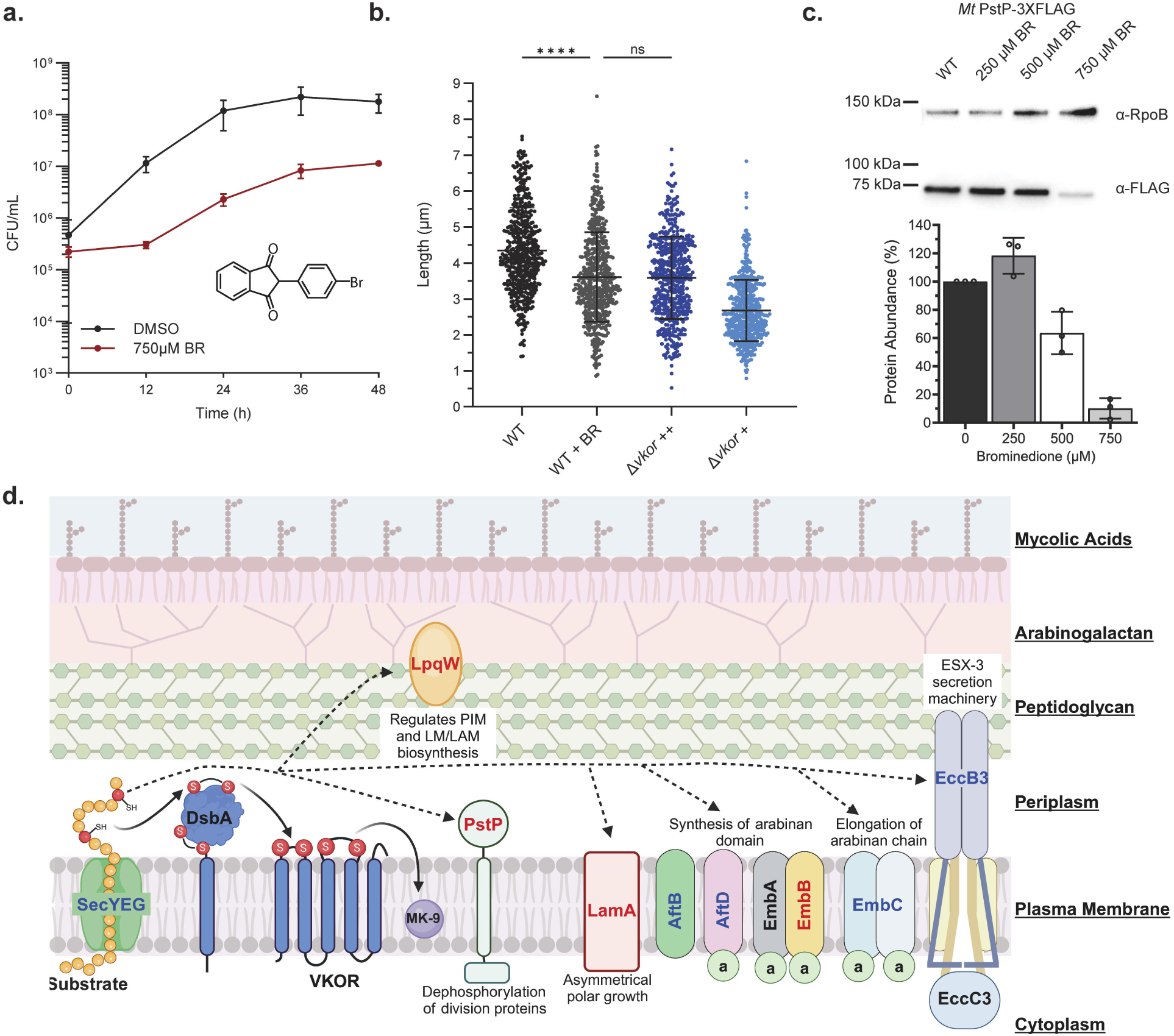
Targeting oxidative protein folding simultaneously affects multiple essential cell envelope proteins. **a,** *M. smegmatis* WT (with DMSO) or treated with 750 µM bromindione (BR) were grown in modified 7H9 broth at 37°C. Data represents the average ±SD of at least two independent experiments. **b,** *M. smegmatis* WT (with DMSO) or treated with 750 µM BR were grown in modified 7H9 broth at 37°C for 36 h. Cells were fluorescently stained with 50 nM Syto24 to stain nucleic acids and 0.6 µg/mL FM4-64 to stain the membrane. Cell dimensions were measured using FIJI (https://fiji.sc/). Data represents the average ±SD. Cell counts included: WT (n=495), WT+BR (n=490), Δ*vkor*++ (n=498), Δ*vkor*+ (n=500). Statistical tests were done using Kruskal-Wallis multiple comparisons. p-values are depicted in GP style: ≤0.0001 (****), 0.0002 (***), 0.021 (**), 0.0332 (*), and non-significant (ns). **c,** *Mt* PstP is unstable when *M. smegmatis* is grown with a VKOR inhibitor (BR). *M. smegmatis* expressing FLAG-tagged proteins were grown with different concentrations of BR at 37°C for 36 h. Proteins were precipitated from cell lysates and reduced before being separated by SDS-PAGE. Protein abundance was determined using α-RpoB as a loading control. Values represent the average ±SD of three independent experiments. **d,** DSBs are present in proteins involved in *M. tuberculosis* cell envelope biogenesis and cell division. See text for further details. Proteins in red text were experimentally demonstrated to have DSBs in this work, while proteins in blue text are predicted substrates identified in this work and structural studies (see Supplementary Table 4). Solid black arrows represent the flow of electrons and dotted black arrows represent the substrates of DsbA/VKOR pathway. MK-9: Menaquinone.

## Discussion

We sought to identify essential client proteins of oxidative protein folding driven by DsbA-VKOR in mycobacteria (Fig. 5d and Supplementary Table 4). We found that 298 out of 4,021 identified proteins are predicted to be secreted and decrease when Δ*vkor* is grown under semipermissive conditions. Additionally, cysteine profiling proteomics identified 381 secreted proteins with oxidized cysteines, from which 132 proteins overlap with downregulated proteins. Together, these findings suggest that ∼14% of the proteome consists of potential substrates of DsbA-VKOR. We showed that the inner membrane proteins EmbB, PstP, and LamA, together with the lipoprotein LpqW require one or two intramolecular DSB(s) for stability. We provide evidence that the two DSBs in *M. tuberculosis* PstP are consecutive and required for survival. We think that the essentiality of DsbA-VKOR may be due to the simultaneous degradation of several misfolded essential proteins.

In addition to these disulfide-bonded substrates, one other protein, EspA, has been found to require an intermolecular DSB^49^. *M. tuberculosis* EspA harbors only one cysteine that allows the formation of a homodimer in both *M. tuberculosis* and *M.smegmatis*^49^. EspA was found in *M. tuberculosis* cell pellets^49^, so it may be translocated to the periplasm where DsbA introduces the DSB to then be secreted as a homodimer. Other examples of disulfide-bonded exported proteins in *M. tuberculosis* have been identified by determining their protein structure (Supplementary Table 4). In these studies, heterologous expression of antigen 85A^50^, LpqW^51^, Culp6^52^, EmbC^53^, MycP1^54^, MycP3^55^, PbpB^56^, AftD^57^, and MmpS5^58^ indicate the presence of DSB(s) when expressed in *E. coli* as well as EccB3^59^ when expressed in *M. smegmatis*. While the presence of DSBs based on structure is an indication of this structural component, for some proteins it is more challenging to determine whether DSBs are present in the native organism. For instance, MycP3 was correctly produced in *E. coli’*s periplasm using a fusion to the *pelB* signal sequence and purified without reducing agents^55^. However, LpqW, Culp6, MycP1, PbpB and EmbC’s carboxy-terminal domain were produced without the signal peptide or TM segment hence in the reductive environment of *E. coli’*s cytoplasm^51–54,56^, and LpqW was reduced during purification^51,60^. So most likely the DSBs formed by air oxidation^61^. On the contrary, the cryoEM structure of *Mt* EmbB does not indicate the presence of DSBs even though the cysteines are close to each other in the structure^62,63^. These studies do not report the use of a reducing molecule during protein purification from *M. smegmatis*^63^ or *E. coli*^62^. Perhaps the amount of protein expressed is too high for DsbA to oxidize/recognize it, and air oxidation did not introduce the DSBs. Thus, determining DSBs in a protein requires expression in the right compartment of the native or closely related host as well as *in vivo* cysteine labeling to ascertain their presence.

The instability by the lack of DSBs and consequent protein degradation was observed for all, LamA, PstP, LpqW, and EmbB. The reduced forms using differential alkylation were also not observed for all except LamA, which suggests that the reduced forms are unstable. On the other hand, the cysteine mutant of EspA is slightly degraded but overall more stable^49^, perhaps because EspA’s DSB is inter-rather than intramolecular and hence not needed for protein stability. Something to note is that the extent of proteolysis varied depending on the protein. For instance, LamA is completely degraded in the Δ*vkor* supplemented with high vs. low cystine. However, there were more oxidized forms of PstP, LpqW, and EmbB when Δ*vkor* was grown with high rather than low cystine, either DsbA may be able to recognize these as a priority or the protease senses more the LamA misfolding over the others. Degradation of misfolded proteins in the periplasm of *E. coli* is one of the functions of the protease DegP^64^. In mycobacteria, there are three orthologs of HtrA (for <u>h</u>igh temperature requirement) protease, including the secreted protease *htrA3*/*pepA* and the membrane-anchored proteases *htrA* and *htrA2*/*pepD*^65^. Of these, only HtrA is essential for survival and the amidase Ami3 is one recently identified substrate^65^. None of these appeared upregulated in the proteomic analysis, however, components of the proteasome including ClpX, Mpa, and Bpa appeared upregulated. Determining which periplasmic protease(s) degrade misfolded proteins in DSB formation mutants will require further investigation.

Proteins that harbor two DSBs can acquire two conformations, either consecutive or non-consecutive. For PstP, the differences in stability of the cysteine mutants suggest that these are consecutive DSBs. As for EmbB, the consecutive cysteines appear close in the structure, so harboring most likely consecutive DSBs. The introduction of non-consecutive DSBs in *E. coli* requires an isomerization pathway directed by DsbCD proteins^66,67^. Previous *in vitro* and structural studies have identified *M. tuberculosis* isomerase-like proteins and a probable re-generating partner^68,69^. However, it is unknown whether isomerization takes place in mycobacteria and what proteins would require isomerization of DSBs in the mycobacterial cell envelope.

The essential catalytic domain of PstP lies in the cytoplasm^39^. In contrast, the periplasmic domain is dispensable and its function remains to be identified. We found that the lack of cysteines or the DSB in the non-essential domain of PstP causes protein instability and degradation. This parallels *E. coli* FtsN, which also contains a DSB in the non-essential SPOR (sporulation-related repeat) domain^70^. However, the extent of degradation in FtsN cysteine mutants does not allow *E. coli* survival^70^, while the degradation observed in the PstP cysteine mutants can partially rescue the growth of the knockdown. These differences may be due to the essential domain of FtsN located on the periplasmic side, whereas the essential domain of PstP lies in the cytoplasm.

We point out that the presence of DSBs in several exported proteins could allow mechanistic studies of protein folding and mycomembrane assembly correspondingly to studies in *E. coli* outer membrane protein folding and assembly^71–74^. Finally, our work emphasizes that targeting DSB formation in the mycobacterial cell envelope would simultaneously compromise several essential proteins as observed with a weak mycobacterial VKOR inhibitor. Better DsbA-VKOR inhibitors would have enhanced bartericidal effect and potentially sensitize mycobacteria to antibiotics targeting the cell envelope.

The limitations of our study include the frequent occurrence of suppressor mutations in the Δ*dsbA* strain which restricted the utility of this conditionally lethal strain. The detection of potential DsbA substrates by our proteomics approach was limited by the use of *M. smegmatis in vitro* growth conditions, where only 60% of the proteome was detected. Virulence factors and toxins specific of Mycobacterial pathogens would not be detected in this model organism. The streptavidin enrichment was 49% selective, better enrichment handles are reported in literature but are not commercially available. The cysteine profiling approach was also constrained by the presence of cytoplasmic proteins, better methods to fractionate the mycobacterial cell envelope could enhance the detection of other oxidized exported proteins by reducing peptide competition for streptavidin binding.

## Methods

### Bacterial strain and growth conditions

Bacterial strains used in this study are listed in Supplementary Table 5. *E. coli* DH10β cells (New England Biolabs) were used for plasmid construction and grown at 37°C in NZ broth (shaking in an Innova incubator at 250 rpm) or agar plates with suitable antibiotics. *M. smegmatis* mc^2^155 was grown at 37°C on NZ agar plates or modified Middlebrook 7H9 broth supplemented with 0.5% bovine serum albumin, 0.2% D-glucose, 0.2% glycerol, 0.05% Tween 80, and additional phosphates including 19.38 mM K_2_HPO_4_, 10.12 mM KH_2_PO_4_ to buffer the acidity of L-cystine. The antibiotic concentrations used were hygromycin 200 µg/mL for *E.* coli and 100 µg/mL for *M. smegmatis*; kanamycin 40 µg/mL for *E.* coli and 25 µg/mL for *M. smegmatis*. *M. smegmatis* Δ*vkor* strains were grown in the presence of 1.5 mM L-cystine (Sigma) unless stated otherwise.

### Strain construction

Primers used in this study are listed in Supplementary Table 6. In order to express *M. tuberculosis* proteins into *M. smegmatis,* genes were amplified from the *M. tuberculosis* H37Rv genome and cloned under the *tet* promoter of pTetG plasmid. The vector was generated by amplifying the pTetG plasmid using primers PR173 and PR151 to then insert *M. tuberculosis pstP* gene amplified with primers PR174 and PR154. The fragments were then ligated using NEBuilder® HiFi Assembly (New England Biolabs) to obtain PL105 plasmid. The right insertion was sequenced with primers PR172 and PR230. Construction of subsequent expression plasmids was made by amplifying the vector from *M. tuberculosis* genome with primers PR173 and PR151 using PL105 as template. The inserts were amplified with primer pairs: PR231 and PR249 for PL148, PR372 and PR373 for PL220, and primers PR330 and PR312 for PL265.

To have an alternative inducible system we constructed a plasmid inducible with acetamide. For this, plasmid PL283 was constructed by removing the TetR of PL105 using primers PR476 and PR477 and re-ligating the product with a KLD mix (New England Biolabs). The intermediate vector was then amplified using PR474 with PR475. The minimal acetamidase regulon^75^ was then amplified from pJV126 using primers PR472 and PR473 and ligated via NEBuilder® HiFi Assembly (New England Biolabs) to the intermediate vector to obtain PL286. The plasmid was confirmed by sequencing with primers PR478, PR479, and PR480.

PstP cysteines were mutagenized by site directed mutagenesis (KLD mix, New England Biolabs) using PL105 as template using primers PR222 and PR223 to construct PL275 and PL299; PR224 and PR225 to make PL276 and PL300; PR278 and PR229 for PL209 and PL302; and PR289 with PR290 for PL291 and PL298. The cumulative cysteine knockout mutants were generated through site-directed mutagenesis reactions using the single cysteine mutant plasmid as the template until all five cysteines of PstP were knocked out.

To silence Ms*pstP* expression, we used a dCas9_Sth1_ system optimized for *M. smegmatis*^47^. The system uses a vector that expresses *Streptococcus thermophilus dcas*9 and the sgRNA scaffold under the control of the Tet promoter (TetR). The PAM sequence CGAGAAG with a score of 1 (a.k.a. high silencing strength) was used^47^. The 20 bp *pstP* targeting region (amino acid 5) was cloned into the sgRNA scaffold using aligned primers PR496 and PR497 (heating equimolar concentrations at 95°C and cooling at 25°C) and assembled with BsmBI-digested vector (Golden gate Assembly, New England Biolabs) using T4 DNA ligase, T4 Polynucleotide Kinase (New England Biolabs). The resulting plasmid, PL282, was sequenced with primer PR291 and transformed into *M. smegmatis* selecting for kanamycin resistance to obtain strain LL464. Induction of *dcas9* with 400 nM aTc caused poor *M. smegmatis* growth.

All plasmids were electroporated into *M. smegmatis* by first growing cells to an OD_600_ 0.8-1.0 in NZ broth. Cells were then washed and resuspended in cold 10% glycerol. An aliquot of 100 µL electrocompetent cells was transformed with ∼500 ng of DNA and electroporated in 2 mm electroporation cuvettes using Gene Pulser II with Pulse Controller Plus and Capacitance Extender Plus accessories (Bio-Rad) with settings 2.5 kV, 1000Ω. and 25 µF. Transformants were recovered using 1 mL of modified-7H9 broth and incubated shaking for 2-4 h at 37°C. Cells were then plated on selective NZ agar plates and/or L-cystine. *M. smegmatis* was independently transformed with plasmids PL105, PL146, PL148, PL220, PL265, PL209, PL243, PL249, PL252, PL258, PL275, PL276, PL288, PL289, PL290, PL291, PL282 to generate the following strains: LL148, LL189, LL191, LL343, LL419, LL398, LL399, LL400, LL401, LL402, LL423, LL424, LL468, LL469, LL470, LL471, LL464, respectively. Similarly, *M. smegmatis* Δvkor was independently transformed with plasmids PL105, PL146, PL148, PL220, PL265 to generate strains LL163, LL194, LL196, LL344, LL420, respectively. Finally, LL464 was independently transformed with the following plasmids: PL286, PL298, PL299, PL300, PL301, PL302, PL303, PL304, PL305, PL306, PL307 to generate the following strains: LL466, LL514, LL515, LL516, LL517, LL518, LL519, LL525, LL526, LL527, LL528, respectively.

### Growth assay

*M. smegmatis* WT, Δ*vkor*, and Δ*dsbA* pTetG*dsbA* were grown in 7H9 broth, supplemented with either 1 mM L-cystine for Δ*vkor* or 0.1 µM of aTc (Sigma) for Δ*dsbA*. Cultures were diluted to an OD_600_ of 0.02 (∼10^5^ cells) in fresh 7H9 media supplemented with 1 mM L-cystine or 0.1 µM of aTc when appropriate. Cultures were incubated at 37°C and shaking at 225 rpm for 48 h. Aliquots were taken at each 12 h timepoint, serially diluted with phosphate-buffered saline (PBS) + 0.05% Tween 80, and plated on NZ media with supplements where necessary. Plates were incubated at 37°C for 72 h to enumerate colonies.

For *MspstP* silencing experiments, overnight cultures were diluted to an OD_600_ of 0.01 in 6 mL of 7H9 broth supplemented with 400 nM aTc and/or 25 µM acetamide. Cells were grown at 37°C for 24 h. Aliquots were serially diluted with PBS + 0.05% Tween 80, and plated on NZ media. Plates were incubated at 37°C for 72 h to enumerate colonies.

### Fluorescence microscopy analysis

*M. smegmatis* WT and Δ*vkor* were grown in 7H9 broth at 37°C for 48 h. Overnight cultures were then diluted to an OD_600_ of 0.01 in 6 mL of 7H9 broth, Δ*vkor* was supplemented with either 1 mM (permissive, ++) or 100 µM (semi-permissive, +). Cells were grown at 37°C to an OD_600_ <0.3, washed, and resuspended in PBS with 0.05% Tween 80. For *MspstP* silencing experiments, overnight cultures were diluted to an OD_600_ of 0.01 in 6 mL of 7H9 broth supplemented with 400 nM aTc and/or 25 µM acetamide. Cells were grown at 37°C for 24 h. Nucleic acid dye, SYTO™24 (Invitrogen), and membrane dye FM™4-64 (Invitrogen) were added to diluted cells at a final concentration of 50 nM and 0.6 µg/mL, respectively. Cells were then incubated for 15 min in the dark and spotted onto a 2% agarose pad. A glass coverslip was laid on top of the agarose and imaged on a Nikon Ti2E microscope equipped with Plan Apo 100x/1.4NA phase contrast oil objective and an sCMOS camera. Several pictures of at least three independent experiments were taken and processed using FIJI^76^ to determine cell dimensions.

### Cysteine profiling proteomics

This protocol was adapated from^34^. Briefly, three *M. smegmatis* WT and four Δ*vkor* biological replicates were grown in 7H9 broth at 37°C for 48 h. Cultures were then diluted in 200 mL (Δ*vkor*) or 50 mL (WT) of 7H9 to an OD_600_ of 0.02. The Δ*vkor* samples were supplemented with 0.5 mM cystine (Δ*vkor*+) and incubated for 43 h at 37°C and 250 rpm of orbital shaking. Cells were harvested and resuspended in lysis buffer containing 50 mM NH_4_HCO_3_, 10 mM MgCl_2,_ 1 mM EDTA, 7 M Urea, 10 µM phenyl-methyl-sulfonyl fluoride (PMSF, Thermo Scientific) pH 7.4, and mechanically lysed using a Mini-Beadbeater (BioSpec). Beating cycles were performed five times for 30 seconds with 2 min ice incubation intervals. Samples were then centrifuged at 13,000 rpm for 10 min at 4°C and cell lysates were subjected to acid-quenching precipitation using 10% TCA (Sigma). Samples were then incubated on ice for 20 min. The protein precipitate was pelleted and washed with cold acetone. Protein pellets were dried and solubilized in 1 mL of labeling buffer (100 mM HEPES pH 8.5 containing 8 M urea, 2% SDS, and 1 mM EDTA). The total protein concentration was determined by Pierce™ BCA assay (Thermo Fisher Scientific) and aimed for 1000 µg of total protein. The sample was then split into two tubes.

One half-sample “diferentially alkylated” was resuspended in blocking buffer with 35 mM iodoacetamide (IAM, Sigma) for 2 h at 37°C in the dark to block all unmodified cysteine residues, while the other half sample “global” was treated with 10 mM TCEP for 1 h at room temperature to reduce all cysteines. Proteins in both half-samples were precipitated with 10% TCA and resuspended in 500 µL of labeling buffer. The diferentially alkylated samples were then reduced with 10 mM TCEP for 1 h at room temperature to reduce disulfide bonded cysteines, while the global samples were treated with 35 mM IAM for 2 h at 37°C in the dark. The counteralkylated samples were TCA precipitated and resuspended in 500 µL of labeling buffer with desthiobiotin iodoacetamide (DBIA; CAS No. 2924824-04-2, Santa Cruz Biotechnology). Both diferentially alkylated and global samples were precipitated by methanol and chloroform. Protein pellets were resuspended in 500 µL of 100 mM HEPES pH 8.0 and digested with LysC and trypsin (Thermo Fisher Scientific) for 16 h at room temperature. Global peptides were flash frozen until use and diferentially alkylated peptides were diluted in binding PBS buffer (0.1 M sodium phosphate, 0.15 M sodium chloride, pH 7.4). Pierce High Capacity Streptavidin Agarose beads were equilibrated as per manufacturers instructions (Thermo Fisher Scientific) and incubated with peptides for 4 h at room temperature. Beads were washed four times with 100 mM HEPES pH 8 and 0.05% NP-40, four times with 100 mM HEPES pH 8 and one time with high purity water to then be resuspended in 50 µL 100 mM HEPES pH 8 and stored at 4°C until processing.

### Peptide purification and labeling

Peptide purification, mass spectrometry analysis, bioinformatics, and data evaluation for quantitative proteomics experiments were performed in collaboration with the Indiana University Proteomics Center for Proteome Analysis at the Indiana University School of Medicine similarly to previously published protocols^77,78^. Following enrichment for DBIA modified peptides, peptides were eluted from the beads with 3 x 100 µL washes of 80 % Acetonitrile, 0.1% formic acid (FA) for 20 min at room temperature, 10 min at room temperature and 10 min at 72 °C all at 600 rpm shaking. These elutions were combined and dried by speed vacuum. For global proteomics, approximately 100 µg of peptides were desalted on Waters Sep-Pak® Vac cartridges (Waters™ Cat No: WAT054955) with a wash of 1 mL 0.1% TFA followed by elution in 3x 0.2 mL of 70% acetonitrile 0.1% FA. Peptides were dried by speed vacuum and resuspended 50 mM triethylammonium bicarbonate (TEAB, from 1 M stock). Each sample was then labeled for two hours at room temperature, with 0.5 mg of Tandem Mass Tag Pro reagent (manufactures instructions Thermo Fisher Scientific, TMTpro™ Isobaric Label Reagent Set; Cat No: 44520, lot no. XC34531). TMTprolabels used for global analysis included, WT three replicas: 127N, 128N, 129N and Δ*vkor* four replicas: 131N, 132N, 133N, 134N, respectively. TMTprolabels used for cysteine profiling analysis included, WT three replicas: 126C, 127C, 128C and Δ*vkor* four replicas: 129C, 131C, 132C, 133C, respectively. Samples were checked to ensure >90 % labeling efficiency and then quenched with 0.3 % hydroxylamine (v/v) at room temperature for 15 minutes. Labeled peptides were then mixed and dried by speed vacuum.

### High pH basic fractionation

Half of the combined global sample and all of the DBIA-peptide sample were resuspended in 0.5% TFA and fractionated on a Waters Sep-Pak® Vac cartridge (Waters™ Cat No: WAT054955) with a 1 mL wash of water, 1 mL wash of 5% acetonitrile, 0.1% triethylamine (TEA) followed by elution for the global sample in 8 fractions of 12.5%, 15%, 17.5%, 20%, 22.5%, 25%, 30%, and 70% acetonitrile, all with 0.1% TEA) and 3 fractions for the DBIA peptide sample of 12.5 %, 22.5 % and 70 % Acetonitrile with TEA.

### Nano-LC-MS/MS

Mass spectrometry was performed utilizing an EASY-nLC 1200 HPLC system (SCR: 014993, Thermo Fisher Scientific) coupled to Eclipse™ mass spectrometer with FAIMSpro interface (Thermo Fisher Scientific). Each multiplex was run on a 25 cm Aurora Ultimate TS column (Ion Opticks Cat No: AUR3-25075C18) in a 50 °C column oven with a 180-minute gradient. For each fraction, 2% of the sample was loaded and run at 350 nl/min with a gradient of 8-38%B over 98 minutes; 30-80% B over 10 mins; held at 80% for 2 minutes; and dropping from 80-4% B over the final 5 min (Mobile phases A: 0.1% FA, water; B: 0.1% FA, 80% Acetonitrile (Thermo Fisher Scientific Cat No: LS122500). The mass spectrometer was operated in positive ion mode, default charge state of 2, advanced peak determination on, and lock mass of 445.12003. Three FAIMS CVs were utilized (–45 CV; –55 CV; –65CV and a technical replicate with –40 CV, –50 CV, and –60 CV) each with a cycle time of 1 s and with identical MS and MS2 parameters. Precursor scans (m/z 400-1600) were done with an orbitrap resolution of 120000, RF lens% 30, 50 ms maximum inject time, standard automatic gain control (AGC) target, minimum MS2 intensity threshold of 2.5e4, MIPS mode to peptide, including charges of 2 to 6 for fragmentation with 60 sec dynamic exclusion shared across the cycles excluding isotopes. MS2 scans were performed with a quadrupole isolation window of 0.7 m/z, 34% HCD collision energy, 50000 resolution, 200% AGC target, dynamic maximum IT, fixed first mass of 100 m/z.

### Mass spectrometry data analysis

Resulting RAW files were analyzed in Proteome Discover™ 2.5.0.400 (Thermo Fisher Scientific^79^ with a *M. smegmatis* UniProt reference proteome FASTA (downloaded 080924, 6600 sequences) plus common laboratory contaminants (73 sequences) and the HPV16E6 protein sequence. SEQUEST HT searches were conducted with full trypsin digest, 2 maximum number missed cleavages; precursor mass tolerance of 10 ppm; and a fragment mass tolerance of 0.02 Da. Static modifications used for the search were: 1) TMTpro label on peptide N-termini and 2) TMTpro label on lysine (K). Dynamic modifications used for the search were 1) carbamidomethylation on cysteine (C) residues (2) DBIA probe (+296.185) on cysteine, 3) oxidation on methionine (M) residues, 4) acetylation on protein N-termini, 5) methionine loss on protein N-termini or 6) acetylation with methionine loss on protein N-termini. A maximum of 3 dynamic modifications were allowed per peptide. Percolator False Discovery Rate was set to a strict setting of 0.01 and a relaxed setting of 0.05. IMP-ptm-RS node was used for all modification site localization scores. Values from both unique and razor peptides were used for quantification. In the consensus workflows, peptides were normalized by total peptide amount with no scaling. Unique and razor peptides were used and all peptides were used for protein normalization and roll-up. Quantification methods utilized TMTpro isotopic impurity levels available from Thermo Fisher Scientific. Reporter ion quantification filters were set to an average S/N threshold of 5 and co-isolation threshold of 30%. Resulting grouped abundance values for each sample type, abundance ratio values; and respective p-values (Protein Abundance based with ANOVA individual protein based) from Proteome Discover were exported to Microsoft Excel. In the global analysis, 4,024 unique proteins were identified, as well as 39,789 peptides groups and 274,054 PSMs. Cysteine profiling analysis identified 13,157 peptides, of which 6,328 were DB-modified peptides (49% enrichment selectivity for cysteine-containing peptides). From these, 5,607 were unique cysteines that belonged to 2546 unique proteins. Secreted proteins containing either signal sequences or transmembrane segments were found using DeepTMHMM 1.0 (https://services.healthtech.dtu.dk/services/DeepTMHMM-1.0/)^80^

### In vivo differential alkylation

*M. smegmatis* WT and Δ*vkor* strains were grown in 7H9 broth at 37°C for 48 h. Cultures were then diluted in 7H9 to an OD_600_ of 0.02. The Δ*vkor* samples were supplemented with 1 mM (permissive, ++) or 0.4-0.5 mM (semi-permissive, +) cystine. Strains expressing FLAG-tagged proteins were induced with 5 nM (EmbB) or 0.2 µM aTc and grown at 37°C for 36 h (PstP and LpqW). LamA-harboring strains, were incubated only 18 h at 37°C with 0.2 µM aTc. Cells were harvested and resuspended in lysis buffer containing 50 mM NH_4_HCO_3_, 10 mM MgCl_2,_ 1 mM EDTA, 7 M Urea, 10 µM phenyl-methyl-sulfonyl fluoride (PMSF, Thermo Fisher Scientific) pH 7.4, and mechanically lysed using a Mini-Beadbeater (BioSpec). Beating cycles were performed five times for 30 seconds with 2 min ice incubation intervals. Samples were then centrifuged at 13,000 rpm for 10 min and cell lysates were subjected to acid-quenching precipitation using 10% trichloroacetic acid (TCA, Sigma). Samples were then incubated on ice for 30 min. The protein precipitate was pelleted and washed with cold acetone twice. Protein pellets were dried and solubilized in 100 mM Tris pH 8 containing 1% SDS. The total protein concentration was determined by Pierce™ BCA assay (Thermo Fisher Scientific) prior to the addition of reducing or alkylating agents.

For *in vivo* differential alkylation, the experimental samples were first treated with 20 mM NEM (N-ethylmaleimide Sigma) and incubated at 37°C for an hour to block free thiols. M63 0.2% glucose media was added to bring the volume to 900 µL and proteins were re-precipitated with 10% TCA followed by acetone washes. Precipitated proteins were then solubilized in 100 mM Tris pH 8 containing 1% SDS and disulfide bonded cysteines were reduced by incubating with 100 mM DTT (dithiothreitol, Sigma) for 30 minutes at room temperature. Samples then underwent another round of TCA precipitation to be subsequently alkylated with either 12.5 mM MalPEG2k (α-[3-(3-Maleimido-1-oxopropyl)amino]propyl-ω-methoxy, polyoxyethylene, NOF America Corporation) or MalPEG5k and incubated for an hour at 37°C. Controls included samples treated with 100 mM DTT and then alkylated with either 20 mM NEM, 12.5 mM MalPEG2k or MalPEG5k using the same conditions described above.

For *in vivo* alkylation, proteins were precipitated as mentioned above and treated with MalPEG2k for an hour at 37°C. Controls included samples treated with 100 mM DTT and then alkylated with either 20 mM NEM or 12.5 mM MalPEG2k using the same conditions described above.

### Western blot analysis and protein quantification

Protein samples were normalized to 2.5-30 µg of total protein, mixed with 5X-SDS reducing loading buffer (100 mM DTT helped to decrease the background), and boiled for 5 min at 100°C (except for EmbB samples which were incubated at room temperature for 15 min) to proceed to SDS-PAGE. Bio-Rad Mini-PROTEAN®TGX™ 4-20% or 7.5% gels were run for 65 min at 150 V. Proteins were then transferred onto PVDF membrane (Millipore) using Trans-Blot SD Semi Dry Transfer Cell (Bio-Rad) for 50 min at 0.07A. Membranes were then blocked with TBS (Tris Buffered Saline) containing 5% milk and incubated with 1:10,000 dilution of α-FLAG antibody (Sigma) overnight at 4°C. Similarly, 1:10,000 dilution of α-RpoB antibody (Invitrogen) was used as a loading control. After incubation with primary antibodies, the blots were washed three times with TBS containing 0.1% Tween 20 and incubated with corresponding secondary antibody either 1:50,000 dilution of α-Rabbit antibody (Santa Cruz Biotechnology) or 1:25,000 dilution of α-Mouse antibody (Santa Cruz Biotechnology) for 1 h at room temperature. Chemiluminescent substrate (ECL, Bio-Rad) was used to detect proteins using ChemiDoc™ MP Imaging System (Bio-Rad). For *in vivo* differential alkylation, proteins were not normalized to load because *vkor* samples would show low signal, thus these were loaded in 5-6 fold excess to be able to see alkylated bands. To determine protein abundance, proteins were treated with DTT only, separated, and transferred to membranes which were cut to blot independently with the two antibodies. The adjusted total band volume was determined for FLAG and RpoB images using Image Lab 6.1 Software. The arbitrary units obtained for each band were then normalized to the volume obtained in the WT protein band. The relative band volume of FLAG was then divided by the relative band volume of RpoB to obtain the relative protein abundance. Anti-FLAG antibody was validated using a total protein extract of *M. smegmatis* without FLAG tag, there were no non-specific bands for α-FLAG raised in rabbit, while α-FLAG raised in mouse produced a non-specific band at ∼50 kDa, which was indicated when appropriate.

### Drug assay

*M. smegmatis* overnight cultures were diluted to an OD_600_ of 0.02 in 6 mL of 7H9 broth supplemented with 750 µM brominedione (CAS1146-98-1, Santa Cruz Biotechnology) and incubated shaking at 225 rpm and 37°C for 48 h. Aliquots were taken every 12 h, serially diluted with PBS + 0.05% Tween 80, and plated on NZ media. Plates were incubated at 37°C for 72 h to enumerate colonies. At 36 h of growth, one aliquot was used to stain with SYTO™24 and FM™4-64 and analyzed by microscopy as described above.

*M. smegmatis* expressing FLAG-tagged *Mt* PstP were inoculated in 7H9 broth to an OD_600_ of 0.02 in fresh 7H9 media containing 0.2 µM aTc and varying concentrations of brominedione (0, 250, 500, and 750 µM) or DMSO (Sigma) as control. Cells were grown at 37°C for 36 h. Proteins were precipitated and reduced to determine abundance using anti-FLAG antibody as indicated above.

### Bioinformatic analysis

*In-silico* analysis was performed on all the previously identified essential genes in *M. tuberculosis*^30^. GI accession numbers of 625 essential genes were first acquired through the Database of essential genes (DEG) database (http://origin.tubic.org/deg/public/index.php) and used to obtain the FASTA protein sequences using the National Center for Biotechnology Information (NCBI, https://www.ncbi.nlm.nih.gov/). To determine which proteins are predicted to be secreted, we used SignalP-5.0^81^ and TOPCONS^82^, which predict the presence of signal sequences and TM segments, respectively. Out of the list of 625 proteins, 47 were predicted to have TM segments and 130 were predicted to be secreted. Proteins were further narrowed down by a more recent essentiality study^36^ to 90 proteins. Through manual curation of TOPCONS plots, proteins predicted to harbor cysteine residues in the extracellular compartment were chosen from the pool, which resulted in a final list of 25 proteins as candidate DsbA-substrates. Finally, a literature search was performed on those to disqualify proteins that were not experimentally demonstrated to be periplasmic^33^ ending with a final list of 19 proteins (Supplementary Table 1).

### Statistical analysis

Comparisons of the cell size mean between mutant and wildtype were done using Kruskal Wallis multiple comparisons with GraphPad Prism version 10.0.0 (USA). Comparisons of the peptide abundance mean between mutant and wildtype were done using non-paired Kruskal Wallis with Dunn’s multiple comparison test using GraphPad Prism version 10.4.1. Comparisons between the survival mean of the wildtype control vs. PstP cysteine mutants were done using ordinary non-paired one-way ANOVA multiple comparisons selecting for Dunnett’s test with GraphPad Prism version 10.4.0. Significant differences were indicated in graphs using GP style: p-value ≤0.0001 (****), 0.0002 (***), 0.021 (**), 0.0332 (*). Non-significant p-values (>0.1234) were indicated as ns.

## Acknowledgements

We appreciate Eranthie Weerapana (Boston College) for helpful advice and discussion while adapting a cysteine profiling protocol to mycobacteria; Julia van Kessel (IU) for her kind gift of pJV126; Xindan Wang and Dan Kearns labs (IU) for their help in microscopy training; Shoko Wakabayashi (Eric Rubin’s lab, HMS) for her help in developing an *in vivo* alkylation protocol in *M. tuberculosis*; Cara Boutte (UTA) for sharing reagents and helpful discussion of PstP experiments; Hesper Rego and Celena Gwin (Yale) for helpful discussion and sharing LamA unpublished data. The mass spectrometry work performed in this work was done by the Indiana University School of Medicine Proteomics Core. Acquisition of the IUSM Proteomics core instrumentation used for this project was provided by the Indiana University Precision Health Initiative. The proteomics work was supported, in part, by the Indiana Clinical and Translational Sciences Institute (funded in part by Award Number UL1TR002529 from the National Institutes of Health, National Center for Advancing Translational Sciences, Clinical and Translational Sciences Award) and, in part, by the IU Simon Comprehensive Cancer Center Support Grant (Award Number P30CA082709 from the National Cancer Institute). Panels in Fig. 1, 3, and 4 were created using Biorender.com with publication licenses IX27IM10UB, HM27JCYKT9, and FW27IM0NXL, respectively. This work was supported by Indiana University Bloomington and by the Cystic Fibrosis Foundation Pilot and Feasibility Award 004846I222 (to C.L.).

## Author’s contributions

Conceptualization, C.L.; Methodology, C.L.; Validation, A.M.S.; Formal analysis, A.M.S. E.D., and C.L.; Investigation, A.M.S., R.C., E.D., and C. L.; Writing – Original Draft, A.M.S., and C.L.; Writing – Review & Editing, C.L.; Visualization, A.M.S.; Supervision and Project Administration, C.L.; Funding Acquisition, C.L.

## Data Availability Section

All data generated or analyzed during this study are included in this published article, its supplementary information, public repository and source data files.

## Source Data

Supplementary Information contains 5 supplementary tables in a word document, Supplementary Table 2 is provided in excel as well as the source data file containing unprocessed images and the data used to generate graphs. Raw and processed mass spectrometry data are uploaded to the MassIVE repository (https://massive.ucsd.edu/) with accession MSV000096877 and ProteomeXchange PXD059898. Data will be made publicly available upon publication.

