## Supplementary material for "Disulfide bonds are required for cell division, cell envelope biogenesis and antibiotic resistance proteins in mycobacteria": Supplementary Information.docx

Running Title: Essential cell envelope proteins harbor disulfide bonds in Mycobacteria

Adrian Mejia-Santana^1^, Rebecca Collins^1^, Emma H. Doud^2,3^, and Cristina Landeta^1*^

Author’s affiliations:

^1^ Department of Biology. Indiana University. Bloomington, IN USA.

^2^ Biochemistry and Molecular Biology. Indiana University School of Medicine. Indianapolis, IN U.S.A.

^3^ Center for Proteome Analysis; Indiana University School of Medicine. Indianapolis, IN U.S.A.

**Keywords:** disulfide bonds, oxidative protein folding, DsbA, VKOR, substrates, essential proteins, PstP, PP2C, Ser/Thr phosphatase, EmbB, Rv2507, MmpS3, LamA, LpqW, MycP3, EccB3, AftD, AftB, mycobacteria, actinobacteria, mycomembrane.

**Supplementary Figures**

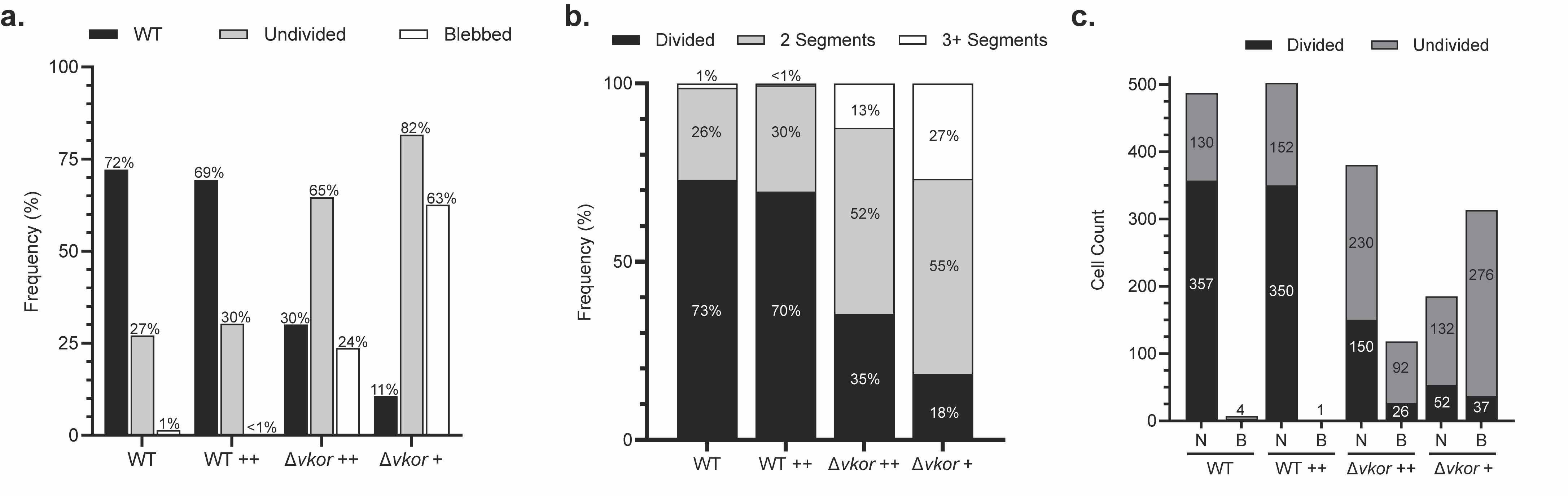
**Supplementary Figure 1.** Lack of DSBs in *M. smegmatis* leads to defects in septation and cell division. *M. smegmatis* WT and Δ*vkor* cells, supplemented with 1 mM (++) or 100 µM (+) cystine, were fluorescently stained with 50 nM Syto24 (nucleic acid stain) and 0.6 µg/mL FM4-64 (membrane stain). **a,** Frequencies of each morphology were obtained using FIJI (https://fiji.sc/) by counting cells with bacillar shape, blebbed, and undivided. Undivided was determined by the presence of FM4-64 stained septum. Some cells displayed more than one phenotype leading to total frequencies above 100%. **b,** Undivided cells were categorized into how many septa were present within one filament and frequencies were obtained relative to the total cell count. **c,** Divided and undivided cells displaying normal bacilli (N) or blebbed (B) morphologies were determined. Cell counts included: WT (n=495), WT++ (n=505), Δ*vkor*++ (n=498), Δ*vkor*++ (n=500).

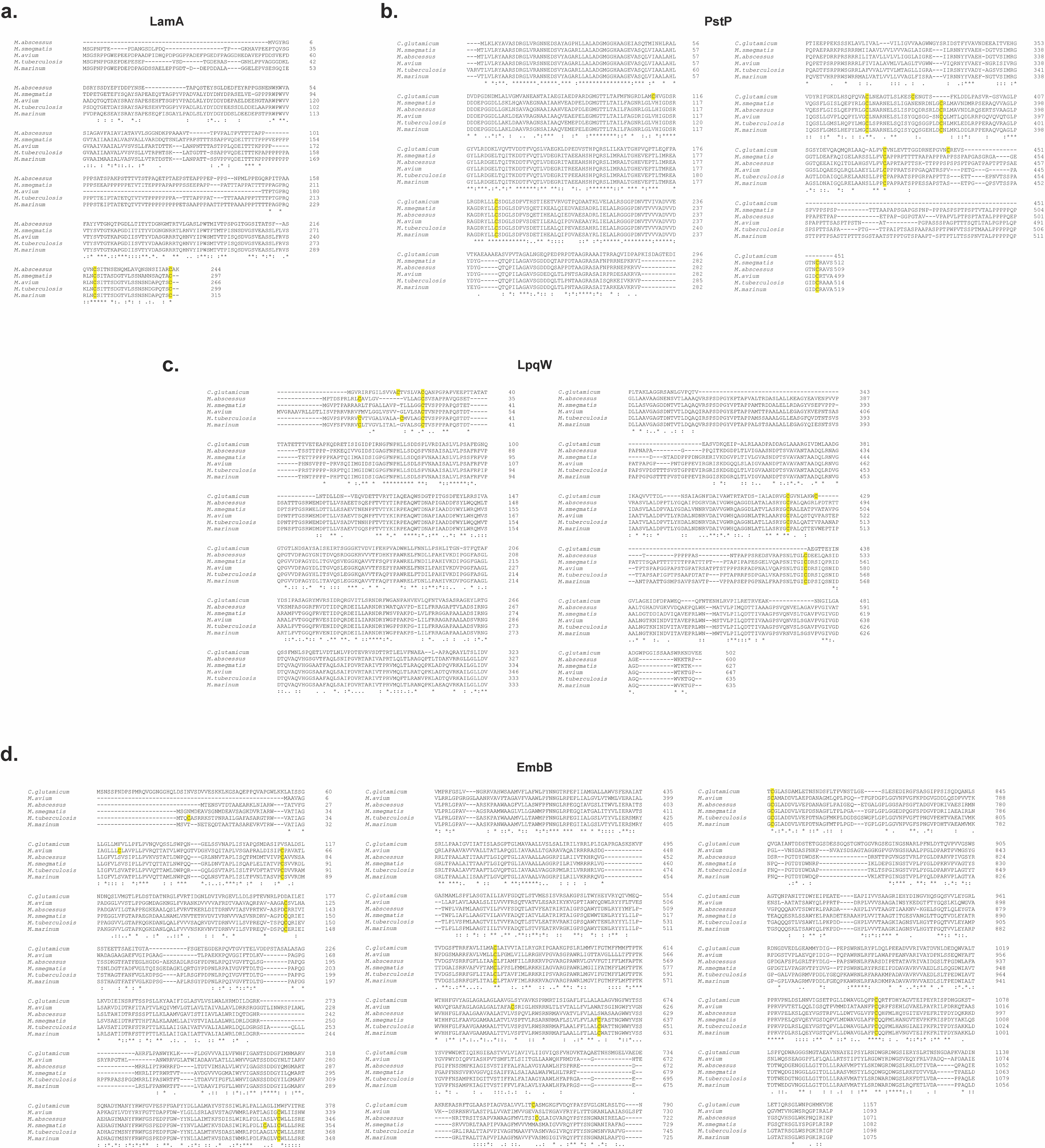
**Supplementary Figure 2.** Cysteine conservation in four predicted *M. tuberculosis* DsbA substrates. Protein sequence alignments of LamA (**a**), PstP (**b**), LpqW, (**c**) and EmbB (**d**) were performed using clustal omega^83^. Reference strains used for the protein sequences include *Mycobacterium avium* 104, *Mycobacterium marinum* M, *Mycobacterium smegmatis* mc^2^155, *Mycobacterium tuberculosis* H37Rv, *Mycobacterium abscessus* ATCC 19977, *Corynebacterium glutamicum* strain R. Cysteines are highlighted in yellow.

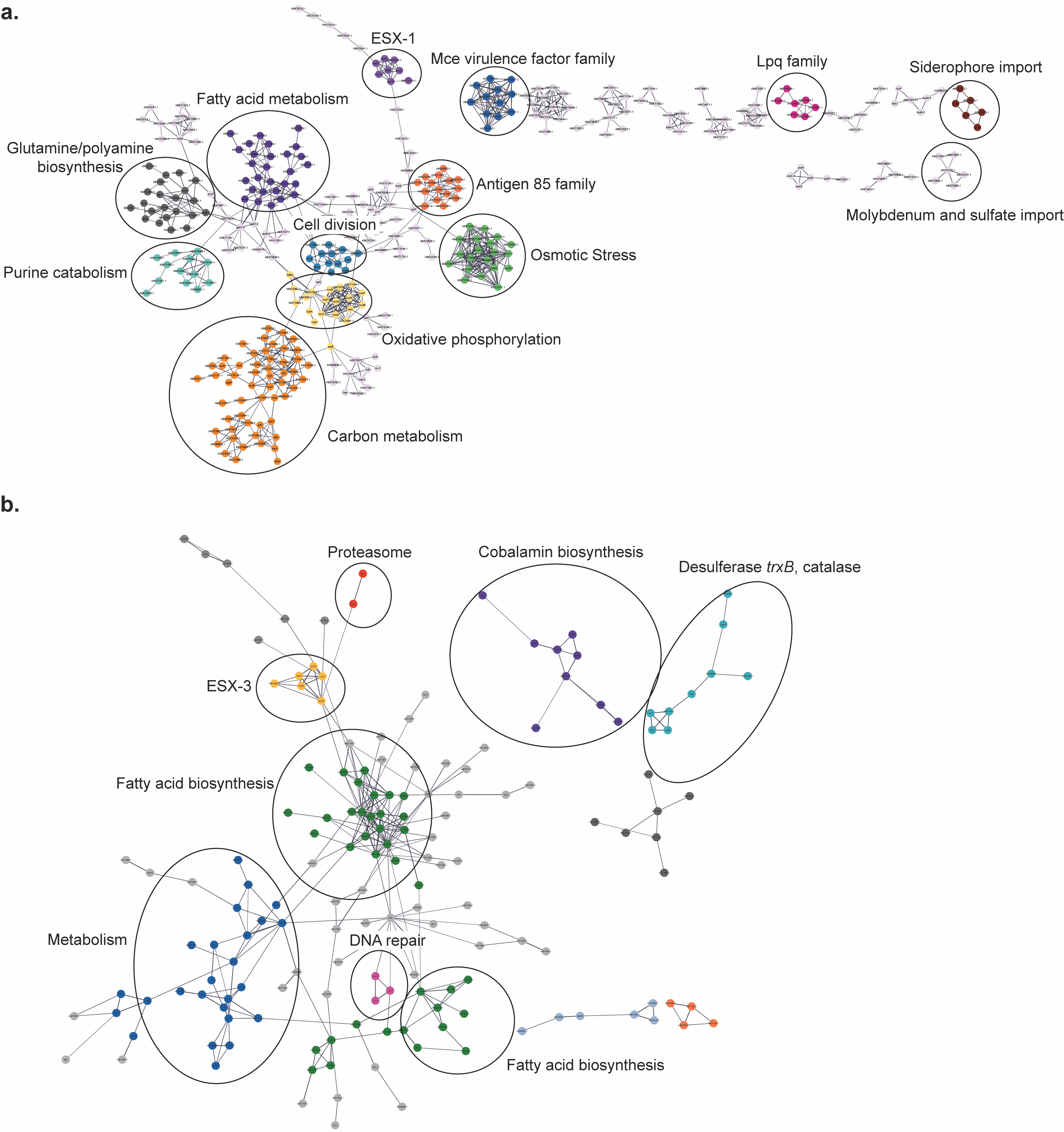
**Supplementary Figure 3.** Protein-protein interactions between (**a**) downregulated and (**b**) upregulated proteins found in Δ*vkor*. Predictions were done using STRING DB V12.0 (https://string-db.org/).

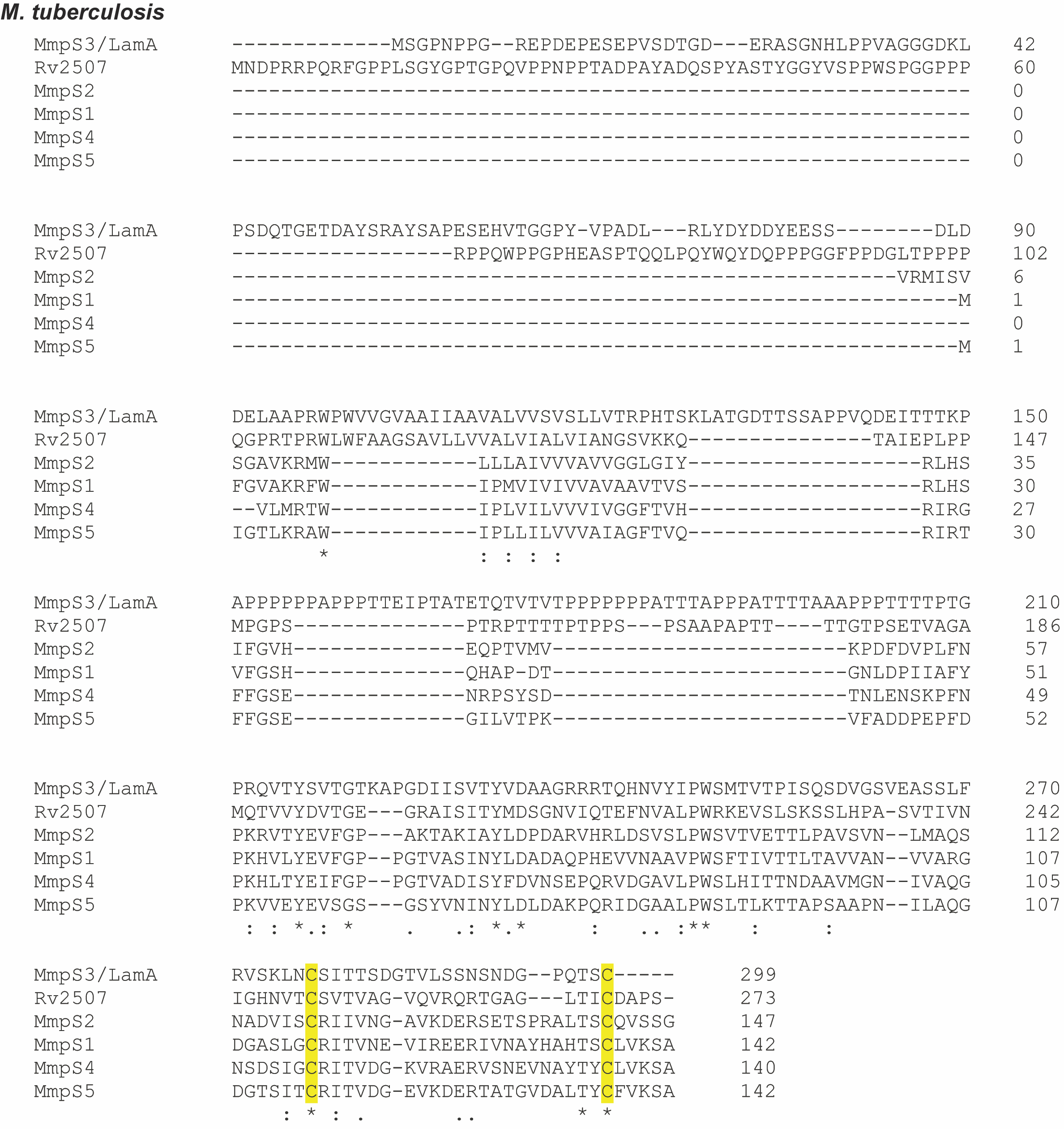
**Supplementary Figure 4.** Cysteine conservation among *M. tuberculosis* H37Rv MmpS paralogs. Protein sequence alignments were done using clustal omega^83^. Cysteines are highlighted in yellow.

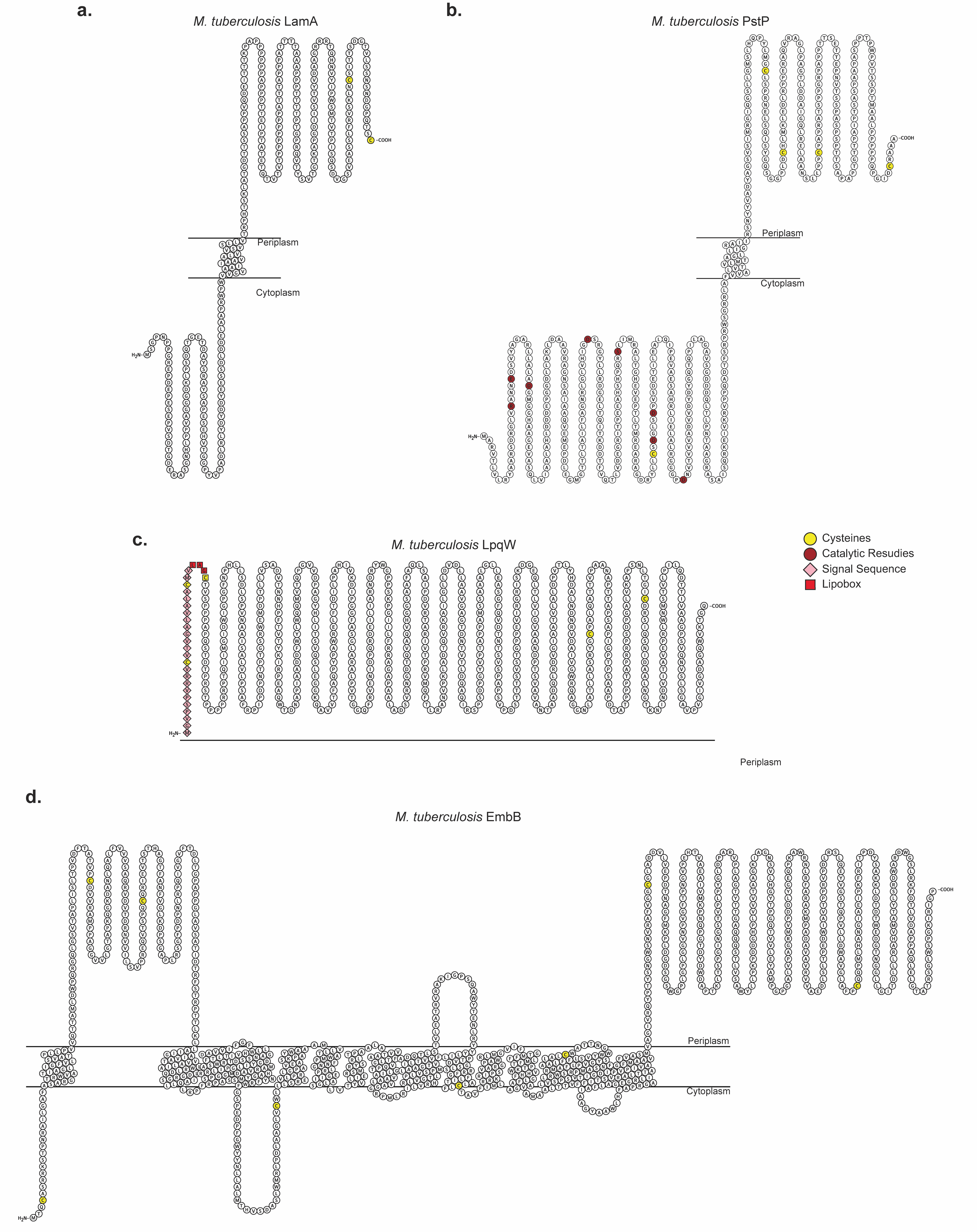
**Supplementary Figure 5.** Membrane protein topologies of four *M. tuberculosis* proteins and the localization of their cysteine residues. Protein topologies of LamA (**a**), PstP (**b**), LpqW (**c**), EmbB (**d**) were visualized using Protter^84^.

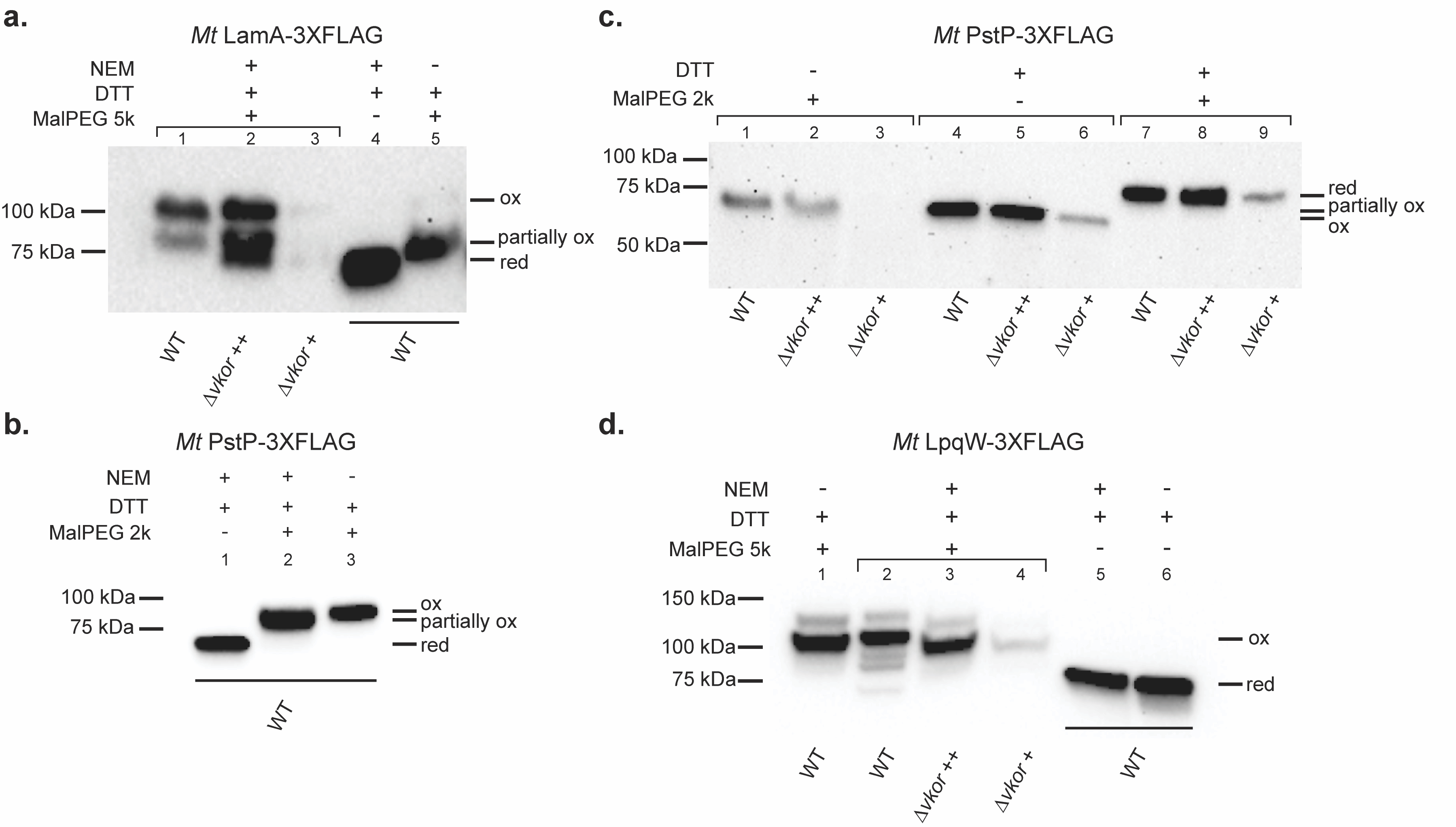
**Supplementary Figure 6.** Differential and conventional alkylation corroborate DSBs in four DsbA substrates. *M. tuberculosis* *Mt*LamA (**a**), *Mt*PstP (**b** and **c**), and *Mt*LpqW (**d**) proteins were fused to a 3X-FLAG tag at their carboxy termini and expressed in *M. smegmatis* WT and Δ*vkor* supplemented with 1 mM (++) or 0.4-0.5 mM cystine (+). Cells were grown at 37ºC in the presence of 200 nM aTc for 36 h (PstP and LpqW) or with 5 nM aTc for 18 h (LamA). Proteins were precipitated from cell extracts and differentially alkylated (**a**, **b** and **d**) by treating them with 20 mM NEM to block free thiols. Disulfide-bonded cysteines were then reduced with 100 mM DTT, and new thiols were alkylated with 12.5 mM MalPEG2k or MalPEG5k when indicated. Controls were treated with 100 mM DTT and then alkylated with either 20 mM NEM or 12.5 mM MalPEG2k or MalPEG5k as indicated. Δ*vkor* samples were loaded in excess to be able to observe alkylated bands. Western blotting using α-FLAG antibody was used to detect the proteins. **c,** *In vivo* alkylation of PstP samples (lanes 1-3) were only alkylated with 12.5 mM MalPEG2k. Controls (lanes 4-9) were treated with 100 mM DTT and then alkylated with 12.5 mM MalPEG2k. Immunoblots are representative images of two independent experiments.

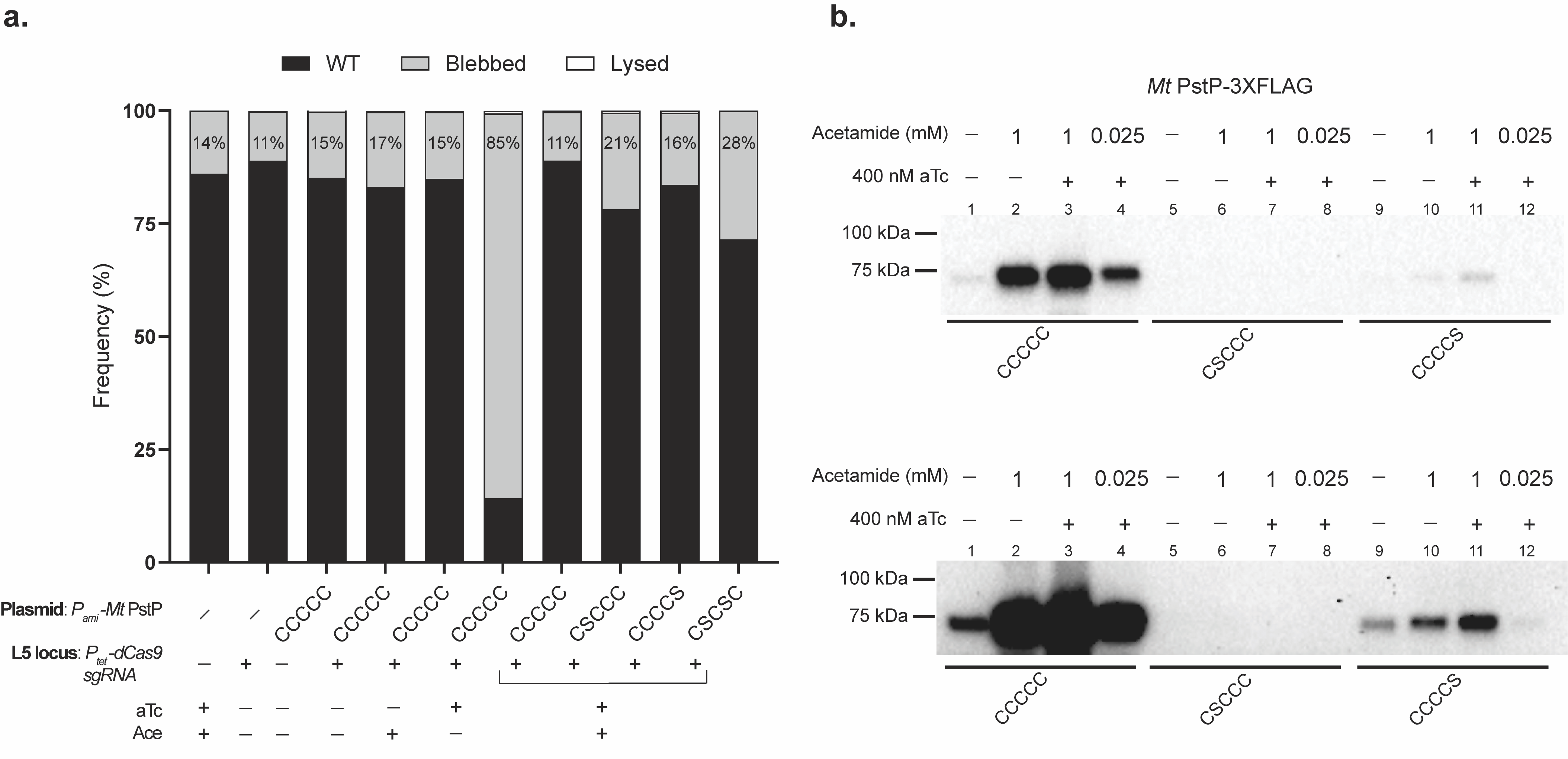
**Supplementary Figure 7.** *M. tuberculosis* PstP cysteine mutants are degraded and display slightly more blebbed events when carrying only one DSB. **a,** *M. smegmatis* *pstP* was silenced using CRISPRi (inducible with aTc), while an ectopic copy of *M. tuberculosis* *pstP*, Cys359Ser or Cys510Ser was used to rescue the knockdown growth. Cells were inoculated to an OD_600_ of 0.01 in 7H9 broth supplemented with 400 nM aTc and/or 25 µM acetamide and incubated at 37ºC for 24 h. Cells were stained and imaged to calculate the frequency of blebbing in 500 cells. **b,** The first DSB between Cys359 and Cys380 of PstP provides more stability than the second DSB between Cys424 and Cys510. Cells were inoculated to an OD_600_ of 0.01 in 7H9 broth supplemented with 400 nM aTc and different concentrations of acetamide. Cells were incubated at 37ºC for 24 h. Proteins were precipitated from cell lysates and reduced with 100 mM DTT. A representative image of two independent experiments is shown. Bottom, overexposed immunoblot to show fainter bands.

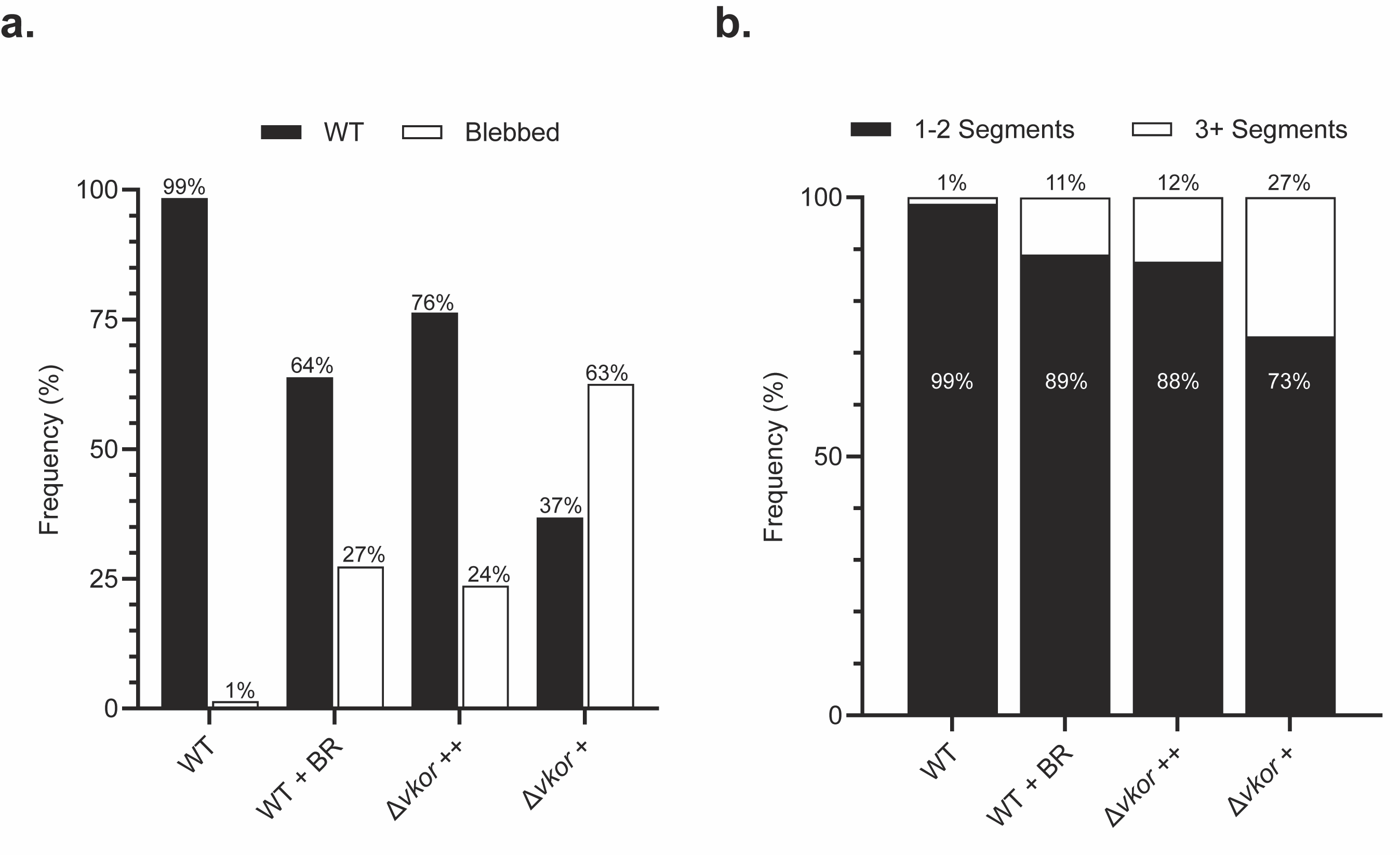
**Supplementary Figure 8.** *M. smegmatis* treatment with a VKOR inhibitor, bromindione (BR), phenocopies the Δ*vkor* morphology grown under high cystine concentration. *M. smegmatis* WT (DMSO control), WT treated with 750 µM of BR, and Δ*vkor* either supplemented with 1 mM (++) or 100 µM (+) cystine, were fluorescently stained with 50 nM Syto24 (nucleic acid stain) and 0.6 µg/mL FM4-64 (membrane stain). **a,** Frequencies of each morphology were obtained using FIJI (https://fiji.sc/) by counting cells with bacillar shape and blebbed. Some cells displayed more than one phenotype leading to total frequencies above 100%. **b,** Undivided cells were categorized into how many septa were present within one filament and frequencies were obtained relative to the total cell count. Cell counts included: WT (n=495), WT+BR (n=490), Δ*vkor*++ (n=498), Δ*vkor*+ (n=500)**.**

**Supplementary Table 1.** Candidate essential substrates of mycobacterial DsbA. *In silico* analysis of 625 essential proteins revealed 19 proteins conserved across all mycobacteria containing extracytoplasmic cysteine residues thus representing candidate substrates of DsbA-VKOR system. GO class was obtained using the gene ontology resource (https://geneontology.org/). #Cys, indicates cysteine residues present in *M. tuberculosis* H37Rv proteins.

|  | **Gene** | ***M. tuberculosis*** | ***M. smegmatis*** | ***M. marinum*** | ***M. avium*** | ***M. abscessus*** | **# Cys** | **GO Class** |
| --- | --- | --- | --- | --- | --- | --- | --- | --- |
| **Cell Envelope Biogenesis/Maintenance** | ***pstP*** | Rv0018c | MSMEG_0033 | MMAR_0020 | MAV_0022 | MAB_0037c | 5 | Serine/threonine protein phosphatase |
|  | ***culp6*** | Rv3802c | MSMEG_6394 | MMAR_5366 | MAV_0216 | MAB_0178 | 4 | Cutinase activity |
|  | ***pbpB*** | Rv2163c | MSMEG_4233 | MMAR_3200 | MAV_2330 | MAB_2000 | 3 | penicillin binding |
|  | ***murX/***  ***mraY*** | Rv2156c | MSMEG_4230 | MMAR_3196 | MAV_2333 | MAB_2003 | 5 | peptidoglycan biosynthesis |
|  | ***embC*** | Rv3793 | MSMEG_6387 | MMAR_5355 | MAV_0225 | MAB_0189c | 7 | Cell wall biogenesis |
|  | ***embB*** | Rv3795 | MSMEG_6389 | MMAR_5357 | MAV_0225 | MAB_0185c | 8 | Cell wall biogenesis |
|  | ***pks13*** | Rv3800c | MSMEG_6392 | MMAR_5364 | MAV_0218 | MAB_0180 | 5 | Lipid metabolism |
|  | ***aftB*** | Rv3805c | MSMEG_6400 | MMAR_5369 | MAV_0212 | MAB_0174 | 5 | Cell wall biogenesis |
|  | ***aftD*** | Rv0236c | MSMEG_0359 | MMAR_0496 | MAV_4923 | MAB_4472 | 7 | Cell wall biogenesis |
|  | ***lpqW*** | Rv1166 | MSMEG_5130 | MMAR_4288 | MAV_1308 | MAB_1315 | 5 | Phospholipid metabolism |
| **Secretion** | ***lepB*** | Rv2903c | MSMEG_2441 | MMAR_1805 | MAV_3758 | MAB_3223c | 6 | Peptidase activity |
|  | ***eccB3*** | Rv0283 | MSMEG_0616 | MMAR_0542 | MAV_4870 | MAB_2233c | 2 | ATPase activity |
|  | ***mycP3*** | Rv0291 | MSMEG_0624 | MMAR_0550 | MAV_4862 | MAB_2225c | 5 | Protease activity |
|  | ***secY*** | Rv0732 | MSMEG_1483 | MMAR_1070 | MAV_4434 | MAB_3784c | 3 | Translocase activity |
| **Uncharacterized** | **Rv1456c** | Rv1456c | MSMEG_3117 | MMAR_2261 | MAV_3324 | MAB_2754 | 2 | heme biosynthetic process |
|  | **Rv0479c** | Rv0479c | MSMEG_0923 | MMAR_0804 | MAV_4671 | MAB_4077 | 2 | Plasma membrane |
|  | **Rv0526** | Rv0526 | MSMEG_0971 | MMAR_0872 | MAV_4619 | MAB_3976c | 3 | Oxidoreductase activity |
|  | **Rv0528** | Rv0528 | MSMEG_0973 | MMAR_0874 | MAV_4617 | MAB_3974c | 3 | Cytochrome complex assembly |
|  | **Rv2507** | Rv2507 | MSMEG_4720 | MMAR_3854 | MAV_1668 | MAB_1539c | 2 | Membrane |

**Supplementary Table 3.** Essential proteins and their overlap found through global and cysteine enrichment proteomic analyses.

| **Proteome wide analysis** | | **Oxidized Cys analysis** | **Essentiality (*M. tuberculosis*)** | ***M tuberculosis* ortholog** |
| --- | --- | --- | --- | --- |
| **Downregulated** | **Upregulated** |  |  |  |
| A0R565 |  | A0R565 | ES | Probable conserved membrane protein |
| A0QUX3 |  | A0QUX3 | ES | lppZ |
| A0QQY8 |  | A0QQY8 | ES | Probable conserved membrane protein |
| A0QPD4 |  | A0QPD4 | ES | aftD |
| A0QNG5 |  | A0QNG5 | ESD | pstP |
|  | A0QQ40 | A0QQ40 | ES | eccC3 |
|  | A0QQ48 | A0QQ48 | ES | eccE3 |
| A0R619 |  |  | ES | Probable conserved membrane protein |
| A0R408 |  |  | ES | prrB |
| A0R205 |  |  | ES | atpE |
| A0R022 |  |  | ESD | pbpB |
| A0R017 |  |  | ESD | ftsW |
| A0QYQ8 |  |  | ES | hypothetical protein |
| A0QNG1 |  |  | ES | pknB |
|  | A0R6H8 |  | ESD | irtA |
|  | A0R5D6 |  | ES | Probable conserved transmembrane protein |
|  | A0R2A3 |  | ES | Probable acyltransferase |
|  | A0QVL8 |  | ES | efpA |
|  | A0QTF1 |  | ES | Probable conserved membrane protein |
|  | A0QQ46 |  | ES | eccD3 |
|  | A0QQ39 |  | ES | eccB3 |
|  |  | A0R2I1 | ES | fdxC |
|  |  | A0R0M4 | ES | ctaD |
|  |  | A0R030 | ES | mptA |
|  |  | A0QWG7 | ESD | Probable conserved membrane protein |
|  |  | A0QVT6 | ES | ftsK |
|  |  | A0QV43 | ES | lepB |
|  |  | A0QV12 | ES | Possible conserved membrane protein |
|  |  | A0QUX0 | ES | gatA |
|  |  | A0QTL8 | ES | aroA |
|  |  | A0QR37 | ES | Probable conserved membrane protein |
|  |  | A0QQB0 | ES | Probable iron-sulfur-binding reductase |
|  |  | A0QP27 | ESD | mmpL3 |
|  |  | A0R7J1 | ES | Probable peptidoglycan hydrolase |
|  |  | A0R7I4 | ES | Hypothetical protein |
|  |  | A0R625 | ES | aftB |
|  |  | A0R614 | ES | embB |
|  |  | A0R613 | ES | embA |
|  |  | A0R612 | ES | embC |

**Supplementary Table 4.** Summary of substrates of mycobacterial DsbA.

| **Protein** | **Gene Locus** | **Essentiality*** | **# Cys** | **DSBs** | **DB-modified Cys^&^** | **Protein production host** | **Ref.** |
| --- | --- | --- | --- | --- | --- | --- | --- |
| Antigen 85A (FbpA) | Rv3804c | NE | 3 | 1 | NA | *E. coli* | ^1^ |
|  | MSMEG_6398 |  | 4 |  | Cys258, Cys281 | *M. smegmatis* | This study |
| LpqW | Rv1166 | ES | 5 | 1 | NA | *E. coli* | ^2^ |
|  | MSMEG_5130 |  | 3 |  | Cys439 | *M. smegmatis* | This study |
| EspA | Rv3616c | NE | 1 | 1 | NA | *M. tuberculosis* and *M. smegmatis* | ^3^ |
| MycP3 | Rv0291 | ES | 5 | 2 | NA | *E. coli* | ^4^ |
|  | MSMEG_0624 |  |  |  | ND | *M. smegmatis* | This study |
| MmpS5 | Rv0677c | NE | 2 | 1 | NA | *E. coli* | ^5^ |
|  | MSMEG_3495 |  |  |  | Cys153 | *M. smegmatis* | This study |
| EmbC | Rv3793 (C’terminal domain) | ES | 7 | 1 | NA | *E. coli* | ^6^ |
|  | MSMEG_6387 |  | 8 | 1 | NA | *M. smegmatis* | ^7^ |
|  |  |  |  |  | Cys138, Cys973 | *M. smegmatis* | This study |
| Culp6 | Rv3802c | ES | 4 | 2 | NA | *E. coli* | ^8^ |
|  | MSMEG_6394 |  |  |  | ND | *M. smegmatis* | This study |
| MycP1 | Rv3883c | NE | 4 | 2 | NA | *E.coli* | ^9^ |
|  | MSMEG_0083 |  |  |  | Cys51, Cys206, Cys244 | *M. smegmatis* | This study |
| PbpB | Rv2163c | ESD | 3 | 1 | NA | *E. coli* | ^10^ |
|  | MSMEG_4233 |  | 2 |  | ND | *M. smegmatis* | This study |
| AftB | MSMEG_6400 | ES | 4 | 1 | Cys357, Cys580, Cys625 | *M. smegmatis* | This study |
| AftD | MAB_4472 | ES | 9 | 2 | NA | *E. coli* | ^11^ |
|  | MSMEG_0359 |  | 7 |  | Cys887, Cys878 | *M. smegmatis* | This study |
| EccB3 | MSMEG_0616 | ES | 3 | 1 | NA | *M. smegmatis* | ^12^ |
|  |  |  |  |  | ND | *M. smegmatis* | This study |
| LamA (MmpS3) | Rv2198c | NE | 2 | 1 | NA | *M. smegmatis* | This study |
|  | MSMEG_4265 |  |  |  | ND |  |  |
| PstP | Rv0018c | ESD | 5 | 2 | NA | *M. smegmatis* | This study |
|  | MSMEG_0033 |  |  |  | Cys186, Cys356, Cys377 |  |  |
| EmbB | Rv3795 | ES | 8 | 2 | NA | *M. smegmatis* | This study |
|  | MSMEG_6389 |  |  |  | Cys85, Cys145 |  |  |

***** NE: Not essential, ES: Essential, ESD: Essential domain^13^.

**^&^** Oxidized cysteine modified with DBIA and identified by mass spectrometry.

ND: Peptides were not detected

NA: Not applicable

**Supplementary Table 5.** List of strains and plasmids used in this study.

| **ID** | **Genotype** | **Ref** |
| --- | --- | --- |
| *Strains* | | |
| NK168 | *M. smegmatis* mc^2^155 | E. Rubin Lab |
| RD149 | *M. smegmatis* mc^2^155 Δ*vkor* | ^14^ |
| NK317 | *M. smegmatis* mc^2^155 Δ*dsbA*::pTetG-*MsdsbA* | ^15^ |
| FLAG-tagged strains | | |
| LL191 | *M. smegmatis* mc^2^155 pTetG-*MtmmpS3*-3xFLAG | This study |
| LL196 | *M. smegmatis* mc^2^155 Δvkor pTetG-*MtmmpS3*-3xFLAG | This study |
| LL148 | *M. smegmatis* mc^2^155 pTetG-*MtpstP*-3xFLAG | This study |
| LL163 | *M. smegmatis* mc^2^155 Δ*vkor* pTetG-*MtpstP*-3xFLAG | This study |
| LL343 | *M. smegmatis* mc^2^155 pTetG-*MtlpqW*-3xFLAG | This study |
| LL344 | *M. smegmatis* mc^2^155 Δvkor pTetG-*MtlpqW*-3xFLAG | This study |
| LL419 | *M. smegmatis* mc^2^155 pTetG-*MtembB*-3xFLAG | This study |
| LL420 | *M. smegmatis* mc^2^155 Δvkor pTetG-*MtembB*-3xFLAG | This study |
| LL189 | *M. smegmatis* mc^2^155 pTetG-*MtmurF*-3xFLAG | This study |
| LL194 | *M. smegmatis* mc^2^155 Δ*vkor* pTetG-*MtmurF*-3xFLAG | This study |
| MtPstP folding mutants | | |
| LL398 | *M. smegmatis* mc^2^155 pTetG-*Mt*pstP_C510S_-3XFLAG | This study |
| LL399 | *M. smegmatis* mc^2^155 pTetG-*Mt*pstP_C510S, C424S_-3XFLAG | This study |
| LL400 | *M. smegmatis* mc^2^155 pTetG-*Mt*pstP_C510S, C424S, C380S_-3XFLAG | This study |
| LL401 | *M. smegmatis* mc^2^155 pTetG-*Mt*pstP_C510S, C424S, C380S, C359S_-3XFLAG | This study |
| LL402 | *M. smegmatis* mc^2^155 pTetG-*Mt*pstP_C510S, C424S, C380S, C359S, C189S_-3XFLAG | This study |
| LL423 | *M. smegmatis* mc^2^155 pTetG-*Mt*pstP_C359S_-3XFLAG | This study |
| LL424 | *M. smegmatis* mc^2^155 pTetG-*Mt*pstP_C380S_-3XFLAG | This study |
| LL468 | *M. smegmatis* mc^2^155 pTetG-*Mt*pstP_C359S, C380S_-3XFLAG | This study |
| LL469 | *M. smegmatis* mc^2^155 pTetG-*Mt*pstP_C359S, C424S_-3XFLAG | This study |
| LL470 | *M. smegmatis* mc^2^155 pTetG-*Mt*pstP_C380S, C510S_-3XFLAG | This study |
| LL471 | *M. smegmatis* mc^2^155 pTetG-*Mt*pstP_C189S_-3XFLAG | This study |
| MsPstP knockdown and cysteine mutants | | |
| LL464 | *M. smegmatis* mc^2^155 L5::P*_tet_-*Sth1-*MspstP* sgRNA, P_tet_-Sth1 dCas9 | This study |
| LL466 | *M. smegmatis* mc^2^155 L5::P*_tet_-*Sth1-*MspstP* sgRNA, P_tet_-Sth1 dCas9, P*ami*-*MtpstP*-3XFLAG | This study |
| LL514 | *M. smegmatis* mc^2^155, L5::P*_tet_-*Sth1-*MspstP* sgRNA, P_tet_-Sth1 dCas9, P*ami*-*MtpstP*_C189S_-3XFLAG | This study |
| LL515 | *M. smegmatis* mc^2^155 L5::P*_tet_-*Sth1-*MspstP* sgRNA, P_tet_-Sth1 dCas9, P*ami*-*MtpstP*_C359S_-3XFLAG | This study |
| LL516 | *M. smegmatis* mc^2^155 L5::P*_tet_-*Sth1-*MspstP* sgRNA, P_tet_-Sth1 dCas9, P*ami*-*MtpstP*_C380S_-3XFLAG | This study |
| LL517 | *M. smegmatis* mc^2^155 L5::P*_tet_-*Sth1-*MspstP* sgRNA, P_tet_-Sth1 dCas9, P*ami*-*MtpstP*_C424S_-3XFLAG | This study |
| LL518 | *M. smegmatis* mc^2^155 L5::P*_tet_-*Sth1-*MspstP* sgRNA, P_tet_-Sth1 dCas9, P*ami*-*MtpstP*_C510S_-3XFLAG | This study |
| LL519 | *M. smegmatis* mc^2^155 L5::P*_tet_-*Sth1-*MspstP* sgRNA, P_tet_-Sth1 dCas9, P*ami*-*MtpstP*_C424S, C510S_-3XFLAG | This study |
| LL525 | *M. smegmatis* mc^2^155 L5::P*_tet_-*Sth1-*MspstP* sgRNA, P_tet_-Sth1 dCas9, P*ami*-*MtpstP*_C359S, C380S_-3XFLAG | This study |
| LL526 | *M. smegmatis* mc^2^155 L5::P*_tet_-*Sth1-*MspstP* sgRNA, P_tet_-Sth1 dCas9, P*ami*-*MtpstP*_C359S, C424S_-3XFLAG | This study |
| LL527 | *M. smegmatis* mc^2^155 L5::P*_tet_-*Sth1-*MspstP* sgRNA, P_tet_-Sth1 dCas9, P*ami*-*MtpstP*_C380S, C510S_-3XFLAG | This study |
| LL528 | *M. smegmatis* mc^2^155 L5::P*_tet_-*Sth1-*MspstP* sgRNA, P_tet_-Sth1 dCas9, P*ami*-*MtpstP*_C359S, C380S, C424S, C510S_-3XFLAG | This study |
| *Plasmids* | | |
| pTetG | Mycobacterial Tet expression vector oriM, colE1 ori, (Hyg^r^) | ^16^ |
| pJR962 | Plasmid for CRISPRi transcriptional repression P*_tet_-*Sth1-sgRNA scaffold, P*_tet_*-Sth1 dCas9, TetR, L5-integrase (Km^r^) | ^17^ |
| pJV126 | Recombineering plasmid with P_hsp60_-*sacB* and acetamidase promoter (Km^r^) | J.van Kessel |
| PL146 | pTetG-*MtmurF-*3XFLAG (Hyg^r^) | This study |
| PL148 | pTetG-*MtlamA*-3XFLAG (Hyg^r^) | This study |
| PL105 | pTetG-*MtpstP-*3XFLAG (Hyg^r^) | This study |
| PL220 | pTetG-*MtlpqW*-3XFLAG (Hyg^r^) | This study |
| PL265 | pTetG-*MtembB*-3XFLAG (Hyg^r^) | This study |
| PL209 | pTetG-*MtpstP*_C510S_-3XFLAG (Hyg^r^) | This study |
| PL243 | pTetG-*MtpstP*_C424S, C510S_-3XFLAG (Hyg^r^) | This study |
| PL249 | pTetG-*MtpstP*_C380S, C424S, C510S_-3XFLAG (Hyg^r^) | This study |
| PL252 | pTetG-*MtpstP*_C359S, C380S, C424S, C510S_-3XFLAG (Hyg^r^) | This study |
| PL258 | pTetG-*MtpstP*_C189S, C359S, C380S, C424S, C510S_-3XFLAG (Hyg^r^) | This study |
| PL275 | pTetG-*MtpstP*_C359S_-3XFLAG (Hyg^r^) | This study |
| PL276 | pTetG-*MtpstP*_C380S_-3XFLAG (Hyg^r^) | This study |
| PL288 | pTetG-*MtpstP*_C359S, C380S_-3XFLAG (Hyg^r^) | This study |
| PL289 | pTetG-*MtpstP*_C359S, C424S_-3XFLAG (Hyg^r^) | This study |
| PL290 | pTetG-*MtpstP*_C380S, C510S_-3XFLAG (Hyg^r^) | This study |
| PL291 | pTetG-*MtpstP*_C189S_-3XFLAG (Hyg^r^) | This study |
| PL282 | pJR962 with *MspstP* (MSMEG_0033) sgRNA (Km^r^) | This study |
| PL283 | pTetG Δ*tetR* (Hyg^R^) | This study |
| PL286 | pTetG Δ*tetR* *amiCA* P*_ami_*-*MtpstP-*3XFLAG (Hyg^r^) | This study |
| PL298 | pTetG Δ*tetR* *amiCA* P*_ami_*-*MtpstP*_C189S_*-*3XFLAG (Hyg^r^) | This study |
| PL299 | pTetG Δ*tetR* *amiCA* P*_ami_*-*MtpstP*_C359S_*-*3XFLAG (Hyg^r^) | This study |
| PL300 | pTetG Δ*tetR* *amiCA* P*_ami_*-*MtpstP*_C380S_*-*3XFLAG (Hyg^r^) | This study |
| PL301 | pTetG Δ*tetR* *amiCA* P*_ami_*-*MtpstP*_C424S_*-*3XFLAG (Hyg^r^) | This study |
| PL302 | pTetG Δ*tetR* *amiCA* P*_ami_*-*MtpstP*_C510S_*-*3XFLAG (Hyg^r^) | This study |
| PL303 | pTetG Δ*tetR* *amiCA* P*_ami_*-*MtpstP*_C424S, C510S_*-*3XFLAG (Hyg^r^) | This study |
| PL304 | pTetG Δ*tetR* *amiCA* P*_ami_*-*MtpstP*_C359S, C380S_*-*3XFLAG (Hyg^r^) | This study |
| PL305 | pTetG Δ*tetR* *amiCA* P*_ami_*-*MtpstP*_C359S, C424S_*-*3XFLAG (Hyg^r^) | This study |
| PL306 | pTetG Δ*tetR* *amiCA* P*_ami_*-*MtpstP*_C380S, C510S_*-*3XFLAG (Hyg^r^) | This study |
| PL307 | pTetG Δ*tetR* *amiCA* P*_ami_*-*MtpstP*_C359S, C380S, C424S, C510S_*-*3XFLAG (Hyg^r^) | This study |

**Supplementary Table 6.** List of primers used in this study

| **ID** | **Sequence 5’-3’** |
| --- | --- |
| PR151 | ggcagcgactacaaagac |
| PR154 | tggtctttgtagtcgctgcctgccgccgcccggcagtc |
| PR172 | cgaggtcgacggtatcgat |
| PR173 | atgtatatctccttcttaattaagcatg |
| PR174 | attaagaaggagatatacatgtggcgcgcgtgaccctgg |
| PR222 | cctgatgggctccctcagcccgc |
| PR223 | taaggctggtgcagggacatgc |
| PR224 | ccctctcgactcccatctgatgaaac |
| PR225 | cccccagactgtccgtag |
| PR226 | gctgccgccttccccggcgccgc |
| PR227 | agggagttggccgccagttcgcgc |
| PR229 | ggctgaggcgggggtggcg |
| PR230 | gtgtacgaccagcacggc |
| PR231 | attaagaaggagatatacatatgagcgggccgaatccc |
| PR249 | tggtctttgtagtcgctgccgcagctcgtttgcggtcc |
| PR278 | gggcatcgactcccgggcggcggcacac |
| PR289 | ttacctgctgtcctcggacgggttgtc |
| PR290 | cgatcaccggcgcgggct |
| PR291 | gctcacatgttctttcctgcgttatcc |
| PR312 | tggtctttgtagtcgctgcctggaccaattcggatcttgcccggtg |
| PR330 | attaagaaggagatatacatatgacacagtgcgcgagcagacgc |
| PR372 | attaagaaggagatatacatatgggcgtgcccagccca |
| PR373 | tggtctttgtagtcgctgccttgcccggtcttcacccattgg |
| PR472 | tcacatgttctttcctgcgaagtgacgcggtctcaag |
| PR473 | ctccttcttaattaagcatgggtcacccctttccattc |
| PR474 | catgcttaattaagaaggagatatac |
| PR475 | acgcaggaaagaacatgtg |
| PR476 | ccatgggctagcggcttt |
| PR477 | gttgcggagccatctagc |
| PR478 | ccaccgctgttctaacgc |
| PR479 | ccagcaacgcggcctt |
| PR480 | gccgcgtatcgcaggac |
| PR493 | tggtctttgtagtcgctgcctgccgccgcccgggagtc |
| PR496 | gggaggtcgctgcgggccgcgtagc |
| PR497 | aaacgctacgcggcccgcagcgacc |
| PR499 | ggcgttcacccttgactt |
